# Phased assembly of neo-sex chromosomes reveals extensive Y degeneration and rapid genome evolution in *Rumex hastatulus*

**DOI:** 10.1101/2023.09.26.559509

**Authors:** Bianca Sacchi, Zoë Humphries, Jana Kružlicová, Markéta Bodláková, Cassandre Pyne, Baharul Choudhury, Yunchen Gong, Václav Bačovský, Roman Hobza, Spencer C.H. Barrett, Stephen I. Wright

**Affiliations:** Department of Ecology and Evolutionary Biology, University of Toronto, Toronto, Canada; Institute of Biophysics of the Czech Academy of Sciences, Brno, Czech Republic; Department of Biology, Queens University, Kingston, Canada; Centre for Analysis of Genome Evolution and Function, University of Toronto, Toronto, Canada; National Centre for Biomolecular Research, Faculty of Science, Masaryk University, Brno, Czech Republic

**Keywords:** *sex chromosomes*, *plants*, *genomics*, *transposable elements*

## Abstract

Y chromosomes are thought to undergo progressive degeneration due to stepwise loss of recombination and subsequent reduction in selection efficiency. However, the timescales and evolutionary forces driving degeneration remain unclear. To investigate the evolution of sex chromosomes on multiple timescales, we generated a high-quality phased genome assembly of the massive older (<10MYA) and neo (<200,000 years) sex chromosomes in the XYY cytotype of the dioecious plant *Rumex hastatulus* and a hermaphroditic outgroup *R. salicifolius*. Our assemblies, supported by fluorescence in situ hybridization, confirmed the neo-sex chromosomes were formed by two key events: an X-autosome fusion and a reciprocal translocation between the homologous autosome and the Y chromosome. The enormous sex-linked regions of the X (296 MB) and two Y chromosomes (503 MB) both evolved from large repeat-rich genomic regions with low recombination; however, the complete loss of recombination on the Y still led to over 30% gene loss and major rearrangements. In the older sex-linked region, there has been a significant increase in transposable element abundance, even into and near genes. In the neo sex-linked regions, we observed evidence of extensive rearrangements without gene degeneration and loss. Overall, we inferred significant degeneration during the first 10 million years of Y chromosome evolution but not on very short timescales. Our results indicate that even when sex chromosomes emerge from repetitive regions of already-low recombination, the complete loss of recombination on the Y chromosome still leads to a substantial increase in repetitive element content and gene degeneration.

## Introduction

One of the most striking and parallel patterns in genome evolution is the degeneration of the non-recombining chromosomes of the heterogametic sex (Y and W chromosomes, hereafter ‘Y’). Sex chromosomes have originated repeatedly across eukaryotes, and while far from universal, signatures of large-scale accumulation of deleterious mutations, the accumulation of repetitive elements and the loss of gene function represent parallel evolutionary outcomes on the non-recombining Y chromosome (Bachtrog 2013; Abbott et al. 2017). Although the extent of degeneration varies greatly among species, many ancient Y chromosomes have lost nearly all their ancestral genes, with evidence of gene retention and sometimes expansion for genes important in reproductive function (Peichel et al. 2020; Subrini and Turner 2021) and meiotic drive (Bachtrog 2020). Despite the widespread recurrent nature of degeneration, our understanding of the timescales over which this occurs, and the evolutionary forces driving Y degeneration remains incomplete.

Several non-mutually exclusive evolutionary processes are thought to contribute to Y degeneration. First, the cessation of recombination causes widespread Hill-Robertson interference between selected sites, weakening the efficacy of natural selection and driving the accumulation of slightly deleterious mutations (Rice 1987; Charlesworth et al. 2005). The loss of recombination can also cause a weakening of selection against transposable elements, both due to Hill-Robertson interference and a reduction in rates of ectopic recombination (Kent et al. 2017). Second, cis-regulatory divergence between the X and Y chromosome can drive loss of gene expression on the Y, enabling a positive feedback loop of expression loss and deleterious mutation accumulation on the non-recombining sex chromosome that can occur even when Hill-Robertson interference effects are weak or absent (Lenormand et al. 2020). Positive selection for gene silencing or loss may also occur on the Y chromosome due to faster rates of adaptation on the X chromosome (Orr and Kim 1998; Crowson et al. 2017) and/or the toxic effects of transposable element activity near genes on the Y (Wei et al. 2020, Muyle et al. 2022). Distinguishing the relative importance of these forces can be challenging, but an improved understanding of the earliest stages of Y degeneration can provide important insights.

The flowering plant *Rumex hastatulus* (Polygonaceae) represents an excellent model system for investigating the timescales and processes driving recombination suppression and Y degeneration. The species has two distinct heteromorphic sex chromosome cytotypes across its geographic range; males to the west of the Mississippi river have one X and one Y chromosome (XY cytotype). Based on our most recent phylogenomic analysis, this sex chromosome system is estimated to have arisen approximately 5-10MYA (Hibbins et al. unpublished data). In contrast, males to the east of the Mississippi have an additional Y chromosome (XYY cytotype), the result of at least one reciprocal translocation event involving the X chromosome and one of the ancestral autosomes (Smith 1964; Kasjaniuk et al. 2019; Rifkin et al. 2021) approximately 180,000 years ago (Beaudry et al. 2020). Our previous work has suggested that the sex-linked regions in this species arose from large tracts of low recombination, particularly in male meiosis, which may have facilitated the evolution of large heteromorphic sex chromosomes (Rifkin et al. 2021, 2022). This includes the neo sex-linked region, which arose from a region of reduced recombination on an ancestral autosome (Rifkin et al. 2021). This system creates an interesting opportunity to study the evolution of sex chromosome regions arising at different (but both young) timescales within the same genetic background.

To better understand the early stages of sex chromosome evolution and Y degeneration, we present a high quality, fully phased assembly of the male genome of the XYY cytotype of *Rumex hastatulus* with highly contiguous assemblies of both Y chromosomes and the fused X chromosome. We characterise the patterns of chromosomal rearrangements, gene loss, and the repetitive DNA accumulation associated with sex chromosome evolution over multiple timescales in this genome. We also sequence and assemble a hermaphroditic species in the genus, *R. salicifolius*, to infer changes in gene order and gene presence/absence evolution on the X and Y chromosomes.

## Results and Discussion

### Genome assemblies

Our phased male genome assembly of the *R. hastatulus* XYY cytotype produced two sets of highly contiguous chromosome-level scaffolds. The ‘maternal’ haplotype had an assembly size of approximately 1,510 MB, with 95% of the genome assembled into four main scaffolds (Figure 1; Tables S1 and S2), which corresponds with the expected chromosome number for the X-bearing haplotype of three autosomes and one sex chromosome (Smith 1964; Rifkin et al. 2021). The BUSCO (Manni et al. 2021) completeness score was 99.3 % (Eukaryota database) and 96.2% (Embryophyta database). Similarly, 97% of the ‘paternal’ assembly was placed into the expected 5 main scaffolds (three autosomes and two Y chromosomes), and an assembly size of 1719 MB, 209 MB larger than the maternal assembly (Figure 1). The BUSCO completeness score was 99.6% (Eukaryota database) and 95.0% (Embryophyta database). The difference in assembly size between the two haplotypes is consistent with previous flow cytometry data, which indicated that the male genome is approximately 10% larger than the female genome (Grabowska-Joachimiak et al. 2015). Cytological measurements suggest the two Y chromosomes combined are approximately 50% larger than the X/NeoX. These findings indicate substantial genome expansion has occurred on the Y chromosomes since they began diverging from the X (see below).

**Figure 1.**
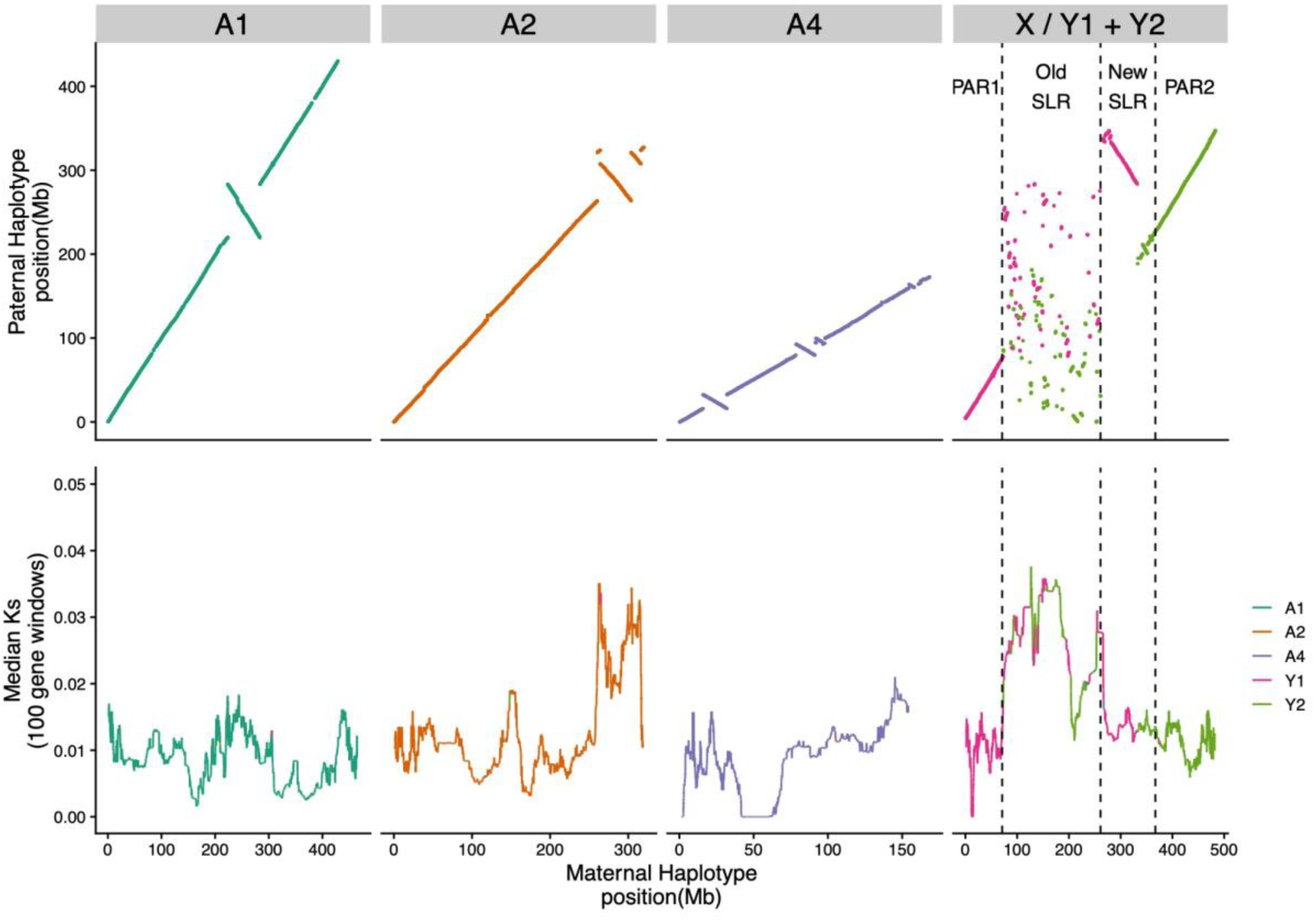
Synteny and divergence between the two assembled haplotypes of XYY male of *R. hastatulus*. Top panel: syntenic genomic position comparison based on whole genome alignment for the autosomes and the sex chromosomes. Bottom panel: median *Ks* between syntenic genes (*Ks*<0.2) along each chromosome, in 100 gene windows with a step size of one gene. Autosomal terminology is used to remain consistent with genome assemblies from the XY cytotype — Autosome 3 (A3) from the XY cytotype is a component of the sex chromosomes in the XYY cytotype. The Old sex-linked-region (‘Old SLR’) is shared with the XY cytotype; PAR1, a pseudoautosomal region, is shared with XY cytotype; the new sex-linked region (‘New SLR’) was formed from sex chromosome fusions; and PAR2, a recombining region, formed from the neo-sex chromosome region.

Our assembly of the hermaphroditic species *R. salicifolius* had a much more compact size of approximately 586MB, with 99.0% of the assembly found in the expected 10 scaffolds, based on chromosome counts of x=10 (Löve 1986). The BUSCO completeness score was 99.6% (Eukaryota database) and 97.1% (Embryophyta database).

Using previously published transcriptome sequences from population samples of both males and females from the XYY cytotype (Hough et al. 2014), we were able to confirm the identification of the sex chromosomes in *R. hastatulus* and validate the high accuracy of the sex-chromosome phasing (Figure S1). In particular, we identified fixed male-specific SNPs and insertion-deletion polymorphisms (indels) from a broad population sample. We found that 7311 out of 7333 fixed sex-specific SNPs and indels mapped to the largest scaffold (hereafter the X chromosome, approx. 483MB) of the maternal haplotype, 7281 (99.3%) of which had the female reference base. Similarly, 99.8% of fixed sex-specific SNPs and indels (6808/6823) mapped to two large scaffolds on the paternal haplotype (hereafter Y1, 343 MB and Y2, 348 MB), and 99.9% of these fixed SNPs and indels contained the male-specific Y base as the reference. These results highlight the high level of completeness and phasing accuracy of the assembled sex chromosomes.

### Synteny analysis

Whole genome alignments integrated with syntenic gene anchors (Song et al. 2022) confirm a high level of synteny across the main autosomes (named according to the naming conventions from the XY cytotype) between the two phased haplotypes of *R. hastatulus* (Figure 1). However, several heterozygous large and small putative inversion differences are apparent across the three main autosomes, indicating a significant degree of inversion heterozygosity. Overall, eight putative heterozygous inversions could be identified on the autosomes, ranging in size from 189 kb to 39 MB in length. These heterozygous inversions collectively span approximately 10% of the autosomes. Strikingly, three of these inversions, including two nested inversions on the second autosome (A2) show highly elevated levels of between-haplotype heterozygosity as measured by *Ks* in gene copies between the haplotypes (Figure 1). Two of these inversions (the nested ones on A2) were independently identified in comparative genetic mapping between the two cytotypes (Rifkin et al. 2021), and these regions as well as the inverted region on A1 were identified as contributing divergent genotype clusters across populations within the XY cytotype (Beaudry et al. 2022). Taken together, these patterns suggest that a subset of these inversion polymorphisms have a deep coalescent time, are shared between the cytotypes and may be subject to balancing selection, potentially due to spatially varying selection, as predicted by theory (Kirkpatrick and Barton 2006) and as observed in several other taxa (Lowry and Willis 2010; Fuller et al. 2019; Todesco et al. 2020; Bieker et al. 2022).

In contrast with the autosomes, a large section of the sex chromosome shows almost no remaining large-scale synteny between the X and Y, highlighting that extensive chromosome rearrangements have occurred since the loss(es) of recombination (Figure 1). Comparisons of the paternal assembly with the previously assembled XY cytotype genome (Figure 2) and patterns of male-specific SNPs from the XY cytotype mapped to the new assembly (Figure S1) reveal that both Y chromosomes contain segments of both the ancestral sex chromosome (‘old sex-linked region’, Figure 1) and much more syntenic segments of the neo-sex chromosomes recently derived from Autosome 3 (‘new sex-linked region’, Figure 1), which recently formed the neo-X and neo-Y chromosome regions. RepeatExplorer (Novák et al. 2013, 2020) analysis and cytogenetic mapping of seven sex-specific satellites (including Cl134 5S together with Cl12 that are located originally on autosome 3 in XY cytotype), in both cytotypes provided further support for the presence of both ancestrally autosomal regions and old sex-linked regions on both Y chromosomes, consistent with our scaffolding results (Figure 3, Figure S3 – S6). Further, the Cl12 and its distribution on the neo-X chromosome suggest that the whole autosome 3 was fused together with the old-X (Figure S3 – S6).

**Figure 2.**
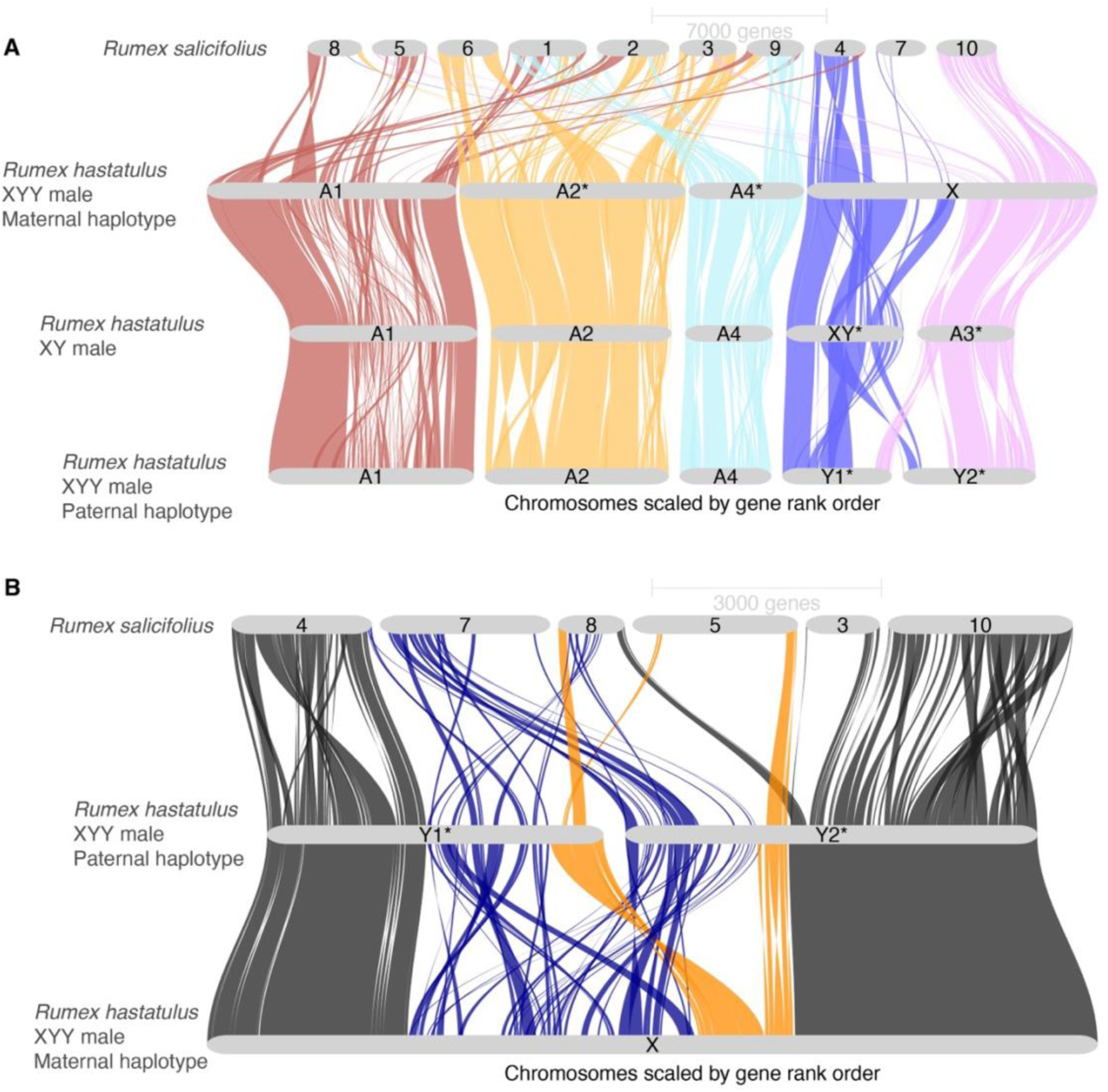
GENESPACE riparian plots between assembled *Rumex* genomes. **A**: Syntenic gene blocks between the *R. hastatulus* XYY paternal and maternal haplotype assemblies, *R. hastatulus* XY male, and *R. salicifolius*. **B**: Close-up view of synteny between the X and Y chromosomes of *R. hastatulus* XYY clade, and orthologous regions in outgroup *R. salicifolius*. The pseudoautosomal regions are coloured in grey, old sex-linked region in blue and new sex-linked region in orange.

**Figure 3.**
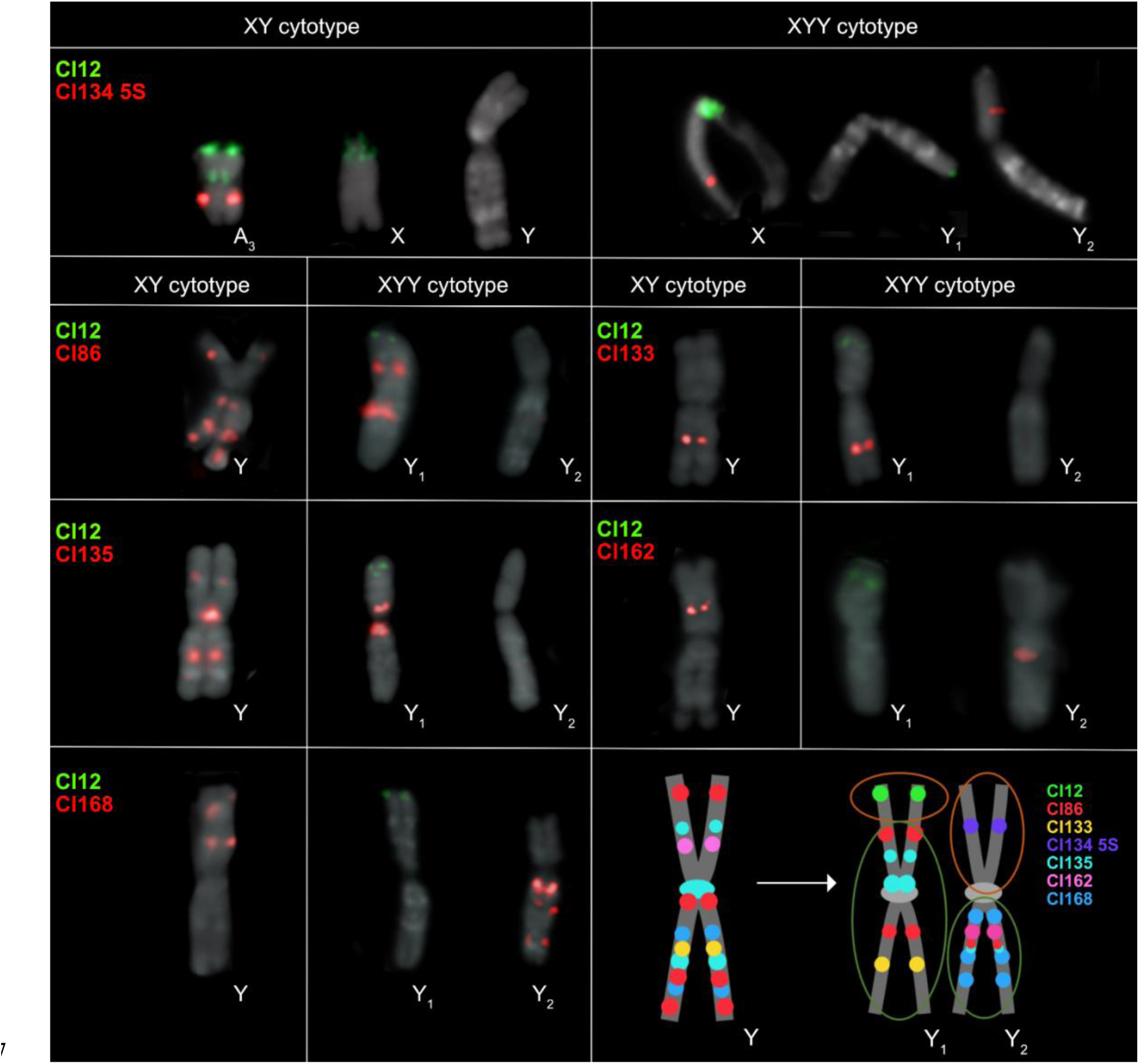
Chromosomal location of seven tandem repeats on the metaphase chromosomes of XY and XYY cytotypes of *R. hastatulus*. The schematic depiction of the structure of the old Y, neo-Y_1_ and neo-Y_2_ chromosomes displays positions of all seven tandem repeats between Y chromosomes in both cytotypes. The position of Cl12 and Cl134 5S changes rapidly in the XYY cytotype, directly confirming translocation and fusion events between autosome 3 (in the XY cytotype) and the X-Y chromosomes. The orange circles represent the parts that originated from the autosome 3 and the green circles represent the old Y-chromosomal regions. Chromosome scale refers to Figure S 3 – 5.

The patterns of fixed sex-linked SNPs from both cytotypes (Figure S1) confirms the presence of a massive sex-linked region (Figure 1), spanning approximately 297 MB on the X chromosome and 503 MB on the Y chromosomes. The absence of sex-limited SNPs at the tips combined with previous comparative genetic mapping results (Rifkin et al. 2021) and early cytogenetic work (Smith 1964) suggest that the sex chromosomes have two pseudoautosomal regions, one on either side of the large, fused X (Figures 1 and 2), where Y1 retains the pseudoautosomal region from the ancestral Y (PAR1), and Y2 contains a pseudoautosomal region derived from the ancestral autosome (PAR2). Altogether these results indicate that, in addition to the X-autosome fusion event, a secondary reciprocal translocation occurred between the homologous autosome and the ancestral Y chromosome. This additional translocation was previously hypothesised from cytological data (Smith 1964) and may have been important to stabilize meiotic pairing, as shown for *Rumex acetosa* Y1XY2 trivalent structure during pachytene synapsis (Cuñado et al. 2007). The difference in outcomes of the reciprocal translocations on the X and Y likely stem from an inversion on the ancestral autosome before or after the fusion, with the X or the translocation with the Ys, as there is no evidence of loss of gene segments on either the Neo-X or the neo-Y segments (Figure 2). This is further supported by FISH results, which show that all main repeat clusters from the ancestral autosome are found on the Neo-X, with evidence of several paracentric and at least one pericentric inversion event on both neo-Ys, further supporting our synteny analysis (Figure 1). Multiple inversion events even between the cytotypes in the old Y-linked regions are evident from the new localisation of satellite clusters Cl86, Cl133, Cl135, Cl162 and Cl168 (Figure 3, S6). It is possible for such large chromosomal rearrangements to occur in a single catastrophic event, as hypothesized in single chromosome shattering in the *Camelina* genome (Mandakova et al. 2019). On the other hand, the satellite enrichment in the XY cytotype could allow for such reorganization, given the new satellite and genome order on the neo-Ys.

In the neo sex-linked regions, synteny is much more retained on this young sex chromosome pair (Figure 1). However, four inversions are apparent within this stretch of approximately 102 MB of new sex-linked sequence, capturing 31% of the region in heterozygous inversions, considerably higher than observed on the autosomes (8 inversions capturing 10% of the sequence in approx. 1GB of the genome). This finding suggests that the recent formation of the neo-sex chromosomes and loss of recombination is accompanied by an elevated maintenance and/or high rate of spread of inversions following the chromosomal fusions.

Comparisons of syntenic gene order in hermaphroditic *R. salicifolius* indicates that, while there have been massive rearrangements genome-wide (Figure 2A), synteny breakdown has been much more extensive on the Y chromosome compared with the X in the sex-linked region (Figure 2B). Specifically, we identify 155 orthologous genes where *R. salicifolius* and the old X chromosome retain syntenic positions whereas the Y position is non-syntenic, and only 13 cases where the old Y and *R. salicifolius* have retained their positions to the exclusion of the X (Table S3). This excess is much greater than the relative difference in non-syntenic orthologues on the autosomes of the two haplotypes (contingency test X^2^ = 26.183, df = 1, *p*<0.001). Interestingly, the pseudoautosomal regions appear to be derived mostly from different ancestral chromosomal origins than the sex-linked regions (Figure 2B). This inference is in line with other chromosomes, where central regions of the chromosome that are associated with large regions of very low recombination (Rifkin et al. 2022) appear to often have been derived ancestrally from different chromosomal regions than the arms, assuming *R. salicifolius* is closer to the ancestral state. The old sex-linked region derives primarily from two *R. salicifolius* chromosomes, scaffolds 7 and 8. To explore whether these two distinct segments represent evolutionary strata that were added to the sex-linked region at distinct times since the formation of the sex-linked region, we estimated *Ks*, the per nucleotide synonymous substitution rate for each sex-linked gene. We found no evidence for a significant difference in the number of ‘young’ (*Ks*<0.03) relative to ‘old’ (*Ks*>0.03) sex-linked genes derived from the two *R. salicifolius* chromosomes (Chi-square contingency test, X^2^= 3.0634, df = 1, *p* = 0.08). Furthermore, while there is heterogeneity across the X chromosome in median X-Y divergence, there is no clear evidence of discrete ‘evolutionary strata’ involving distinct chromosomal segments in the old sex-linked region (Figure 1). This finding may be due to the extensive chromosomal rearrangements that have occurred since the origins of the sex-linked region, the origins of the sex-linked region from a pre-existing region of reduced recombination without strata and/or an ongoing history of gene conversion between some sex-linked genes.

### Genomic distribution of repeats

Previous work indicated that all *R. hastatulus* chromosomes have large, repeat-rich regions of low recombination, including the sex-linked regions (Rifkin et al. 2021, 2022). A resulting question is whether further loss of recombination on the sex-linked regions of the Y chromosomes drives additional and distinct repeat accumulation. Despite the high levels of repetitive content genome-wide (84% and 86% on the maternal and paternal haplotype, respectively), repeat annotation of our phased assemblies reveals that the Y chromosomes have considerably more TEs than the X or autosomes (Figure 4). Mutator-like DNA elements show a major localised accumulation on Y1 and a minor accumulation on Y2, copia-like elements show additional accumulation on Y2, and Ty3 elements have accumulated in localised positions on both Y1 and Y2 (Figure 4, S8). The older sex-linked regions of both Y1 and Y2 have higher repeat content than the older sex-linked regions of the X (Figure 4B). Overall copy number is significantly elevated by almost three-fold on the old sex-linked Y region compared with the old sex-linked X (Table S4, Figure S7B; Chi squared test, *p*<<0.001). In contrast, TE copy number is marginally elevated (1.09-fold) on the newly sex-linked region of the X compared to the Y, only slightly higher than the baseline difference between the sex chromosomes in the PAR (1.02-fold) (Table S4). This likely reflects stochastic differences between haplotypes and/or minor technical differences in TE annotations.

**Figure 4:**
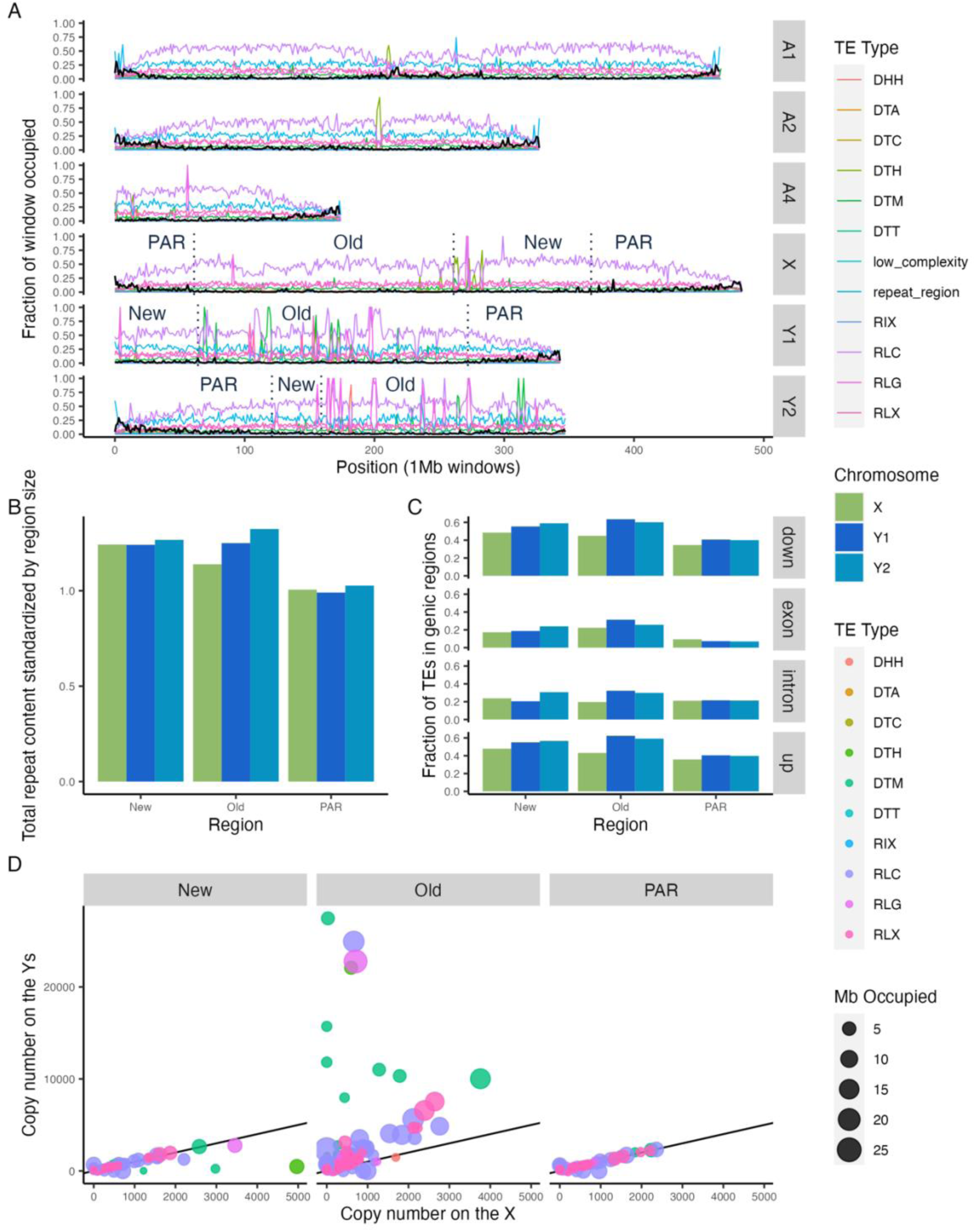
TE distribution across the genome of *Rumex hastatulus* indicates increased TE content on the Y chromosomes relative to the X chromosome or autosomes. **A**: Proportion of 1 MB non-overlapping windows occupied by genes and TE sequences. Values for autosomes and Y chromosomes are from the paternal haplotype, except for the values shown on the X, which are from the maternal haplotype. Black lines are the fraction of the window that is genic sequence and coloured lines are fractions of the window that is TE sequence. The dotted vertical lines and labels on the sex chromosomes divide the chromosomes into newly sex-linked (New), old sex-linked (Old), and pseudo-autosomal regions (PAR). **B**: The summed length of annotated repeats on the newly sex-linked, old sex-linked, and pseudo-autosomal regions of the sex chromosomes divided by the size (total bp) of the region. Considerable nesting and overlap of repeats have led to ratios larger than one. **C**: For each gene on the sex chromosomes, the average proportion of nearby TEs within the newly sex-linked, old sex-linked, and pseudo-autosomal regions. The genic regions were separated into, in order from top to bottom: 1kb upstream, within exons, within introns, and 1kb downstream. The X genes are LiftOff matches from the paternal haplotype and the Y genes were filtered for those that LiftOff found a match for in the maternal haplotype. **D**: Each point represents a family of TEs. The size of the point reflects the number of Mb occupied by members of the family (includes all sex chromosomes) and the X and Y axis values reflect the number of family members on the X and Y chromosomes, respectively. The black line has a slope of 1, to visually indicate where equal quantities would fall. The three-letter transposon codes in the legend are from Wicker et al. 2007. DHH: Helitron, DTA: hAT, DTC: CACTA, DTH: Harbinger, DTM: Mutator, DTT: Tc1-Mariner, RIX: unknown LINE (long interspersed nuclear element), RLC: Copia, RLG: Ty3, RLX: unknown LTR (long terminal repeat).

Transposon families are a useful unit of comparison for understanding TE abundance in the two haplotypes. Wicker et al. (2007) proposed an 80-80-80 rule of similarity to group transposons into families. The procedure requires that the TEs be at least 80bp in length and have 80% similarity over 80% of the aligned sequences. PanEDTA uses this definition to group the annotated TEs into families across the two haplotypes, which allows for a more direct comparison of TE complement. Many individual TE families occupy more space and are more numerous on the older sex-linked regions of the Y chromosomes relative to the X (Figure 4D). This pattern is especially true for harbinger, mutator-like, and LTR elements (Figure S7). Some of this accumulation has led to extreme clusters of very high copy numbers on the old Y, suggestive of local targeted transposition and/or expansion via tandem arrays (Figure S8).

These results suggest extensive accumulation of transposable elements has occurred on the older sex-linked regions of the Y chromosome, but it is unclear whether this accumulation may be affecting genes. To understand whether this TE accumulation is primarily occurring in already repeat-dense areas, the overlap between the TE annotation and gene annotation was examined. To make comparisons as equivalent as possible for this analysis, we used a gene liftover (see Methods) of the paternal genome annotation to the maternal genome annotation and only retained genes with at least one open reading frame in both the original and lifted over annotation.

We observed significantly elevated numbers of TEs inside and near genes on the Y chromosomes, particularly in the old sex-linked region (Figure 4C, Table S4). In contrast, genes in neo sex-linked regions showed no signs of rapid TE accumulation on the Y, as differences between X and Y are similar to baseline differences between the PARs (Figure 4C, Table S4). Overall, we found signs of considerable accumulation of TEs in our older sex-linked region of the Y chromosomes, including into and near genes, although to a lesser extent than TE accumulation further from genes.

### Gene retention and loss

Previous studies of gene loss using transcriptome and short-read genome information on plant sex chromosomes have focused on the pairwise comparison of X and Y chromosomes (Hough et al. 2014; Bergero et al. 2015; Papadopulos et al. 2015; Beaudry et al. 2017; Crowson et al. 2017). This approach cannot distinguish between gene loss and gene movement or duplication among sex chromosomes and autosomes. The genome of a hermaphroditic outgroup, in this case our *R. salicifolius* assembly, allows for the specific identification of genes not present on one of the *R. hastatulus* sex chromosomes that were ‘ancestrally’ present in the same syntenic block. This in turn enables quantification of the extent of *bona fide* gene loss on the sex chromosomes by identifying syntenic orthologs in the outgroup.

Compared to all autosomes and the neo-sex chromosome, there is a high proportion (∼34%) of genes in the old sex-linked region that show evidence of loss on the Y chromosome despite their syntenic presence in both *R. salicifolius* and the X chromosome (Figure 5A, Table S5). Approximately 38% of the lost genes still showed fragments on the Y chromosome, and were classified as partially lost, whereas the remainder are inferred to be fully deleted. These estimates are much higher than on autosomes or the X chromosome, suggesting that the extent of loss is much greater than expected simply from gene copy number variation and/or bioinformatic errors. Overall, if we use the autosomal ‘loss’ values as a baseline for presence-absence polymorphism and/or technical error, we see approximately 30% of genes have been lost on the Y chromosome in the old sex-linked region. Patterns of gene loss along the Y chromosome show evidence of regional variation in the extent of loss, particularly when anchored to the *R. salicifolius* genome with a likely more ancestral gene order (Figures 4B and C). This finding could reflect either the presence of large-scale regional deletions and/or a dynamic history of recombination suppression (i.e. evolutionary strata).

**Figure 5.**
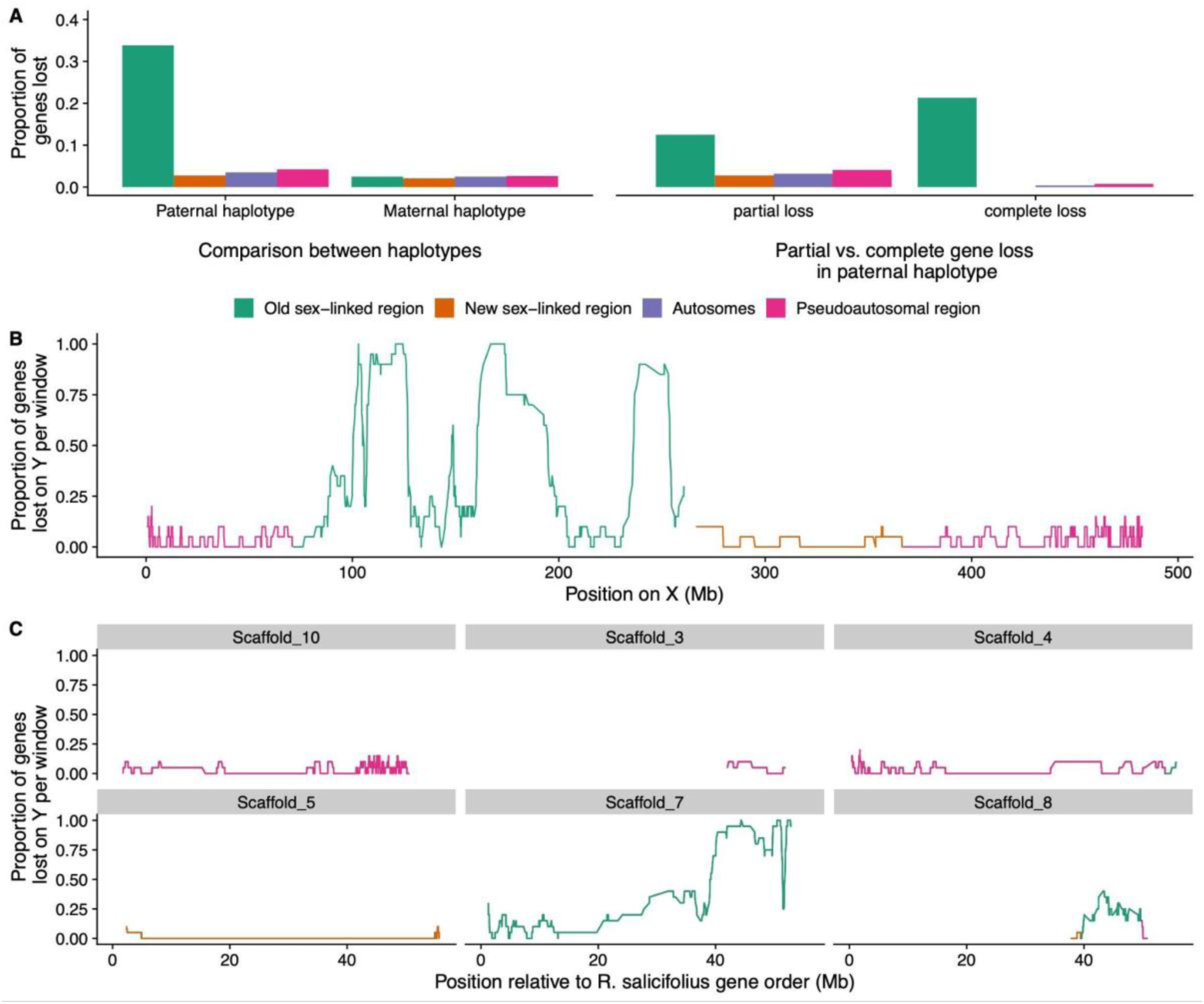
**A**: Proportion of genes lost within the sex-linked regions, pseudoautosomal regions and autosomes in *Rumex hastatulus*. Left panel: comparison of genes which are absent from the paternal (Y-bearing) haplotype while present on maternal (X-bearing) haplotype assemblies, and vice versa. Right panel: proportion of partially lost vs. completely lost genes across regions. **B**: Proportion of genes lost on Y calculated in 20 gene windows along the X chromosome. **C**: Proportion of genes lost on Y calculated in 20 gene windows with respect to the positions of their syntenic genes in *R. salicifolius* scaffolds.

In contrast, we see no sign of excess gene loss on the old X-linked region (Table S6, Figure 5A), providing no evidence of early gene loss on the X chromosome, as found recently in other systems (Mrnjavac et al. 2023). Furthermore, there is no sign of excess gene loss in the ‘new’ sex-linked region (NeoY), suggesting a lack of rapid deletion of Y-linked genes since the chromosomal fusion. Among the genes lost in the neo-X and autosomes, almost all are classified as partially lost. In particular, the evidence for complete gene loss of syntenic orthologs is nearly exclusively restricted to the old Y (159 genes fully lost on the Y, compared with only 24 completely lost in the rest of the genome).

## Conclusions

Our results provide two timepoints early in the evolution of heteromorphic sex chromosomes, supported by a hermaphroditic outgroup. Our studies revealed that in the extremely young (<200,000 generations) neo-sex chromosome of *R. hastatulus*, chromosome rearrangements have accumulated rapidly without signs of gene loss or transposable element invasion. On the older (but relatively young, <10 MYA) regions of the sex chromosomes, extensive rearrangements have led to a near-complete breakdown of synteny, transposable element invasion, and extensive gene loss. The extent of rearrangement is striking for a relatively young sex chromosome system which retains low X-Y divergence for many of the genes that remain. This extent of Y degeneration and sex chromosome evolution is in line with recent results from an unrelated dioecious plant, *Silene latifolia,* which has an approximately 11MYA plant Y chromosome and includes strata as young as 5MYA (Moraga et al. 2023, Akagi et al. 2023). The emergence of sex-linked regions in large pericentromeric regions of low recombination may contribute to a highly dynamic genetic system that evolves heteromorphic sex chromosomes over a relatively short time period.

## Methods

### Long read genome sequencing

We grew a male and female plant from two independent maternal families of *R. hastatulus* from the XYY clade collected from Marion, South Carolina (Pickup and Barrett 2013) in the University of Toronto glasshouse. Following full-sib mating from this F_1_ generation, 11g of leaf tissue from a single F_2_ male was sampled to extract high-molecular weight DNA conducted by Dovetail Genomics (Cantata Bio, LLC, Scotts Valley, CA, USA). 4,618,456 PAC Bio CCS reads (Pacific Biosciences Menlo Park, CA, USA) were sequenced by Dovetail for a total of 87.7 GB (approximately 46x coverage, based on a male genome size estimate of 1.89 GB (Grabowska-Joachimiak et al. 2015). Similarly, we ordered a single *R. Salicifolius* plant from seed collected from Nevada USA from the United States Department of Agriculture’s US National Germplasm System (Accession RUSA-SOS-NV030-372-10), and collected 20 g of leaf tissue for high-molecular weight DNA extraction and sequencing. A total of 5,149,926 PAC Bio CCS reads were sequenced totalling 75.3 GB (approximately 108X coverage based on our flow cytometry estimate of 696 MB).

### Dovetail Omni-C library preparation and sequencing

Proximity ligation and sequencing was conducted by Dovetail using Omni-C sequencing for both species. For each Dovetail Omni-C library, chromatin was fixed in place with formaldehyde in the nucleus and then extracted. Fixed chromatin was digested with DNAse I, chromatin ends were repaired and ligated to a biotinylated bridge adapter followed by proximity ligation of adapter containing ends. After proximity ligation, crosslinks were reversed, and the DNA purified. Purified DNA was treated to remove biotin that was not internal to ligated fragments. Sequencing libraries were generated using NEBNext Ultra enzymes and Illumina-compatible adapters. Biotin-containing fragments were isolated using streptavidin beads before PCR enrichment of each library. The libraries were sequenced on an Illumina HiSeqX platform to produce an approximately 30x sequence coverage.

### De novo assembly

For the male *R. hastatulus* sample, we conducted a haplotype-resolved de novo assembly using Hifiasm (Cheng et al. 2021), using the Omni-C sequencing for haplotype resolution. Paired-end OmniC reads were then mapped and filtered to the two phased assemblies using bwa v0.7.15) (Li and Durbin 2009) following the Arima mapping pipeline (https://github.com/ArimaGenomics/mapping_pipeline), and resulting filtered (MapQ>10) bam files had duplicates marked using Picard v2.7.1. We scaffolded both haplotypes of the assembly using YAHS (Zhou et al. 2023) to generate scaffolded assemblies from each phased haplotype. We manually inspected the scaffolded assembly using a combination of Juicebox (Durand et al. 2016) and whole genome alignment to our previous assembly from the XY cytotype (Rifkin et al. 2022) to identify and break one false join in the assembly. In particular, a break was inserted at the point between autosome 4 and Y2 in haplotype 2 based on manual inspection. Note that we refer to the haplotypes as maternal and paternal based on their sex chromosome composition, but the nature of the phased assembly means the parental origins of each autosome haplotype is not known.

For R. *salicifolius,* Hifiasm was run by Dovetail to generate the primary scaffolds. BLAST (Altschul et al. 1990) results of the *R. salicifolius* Hifiasm output assembly against the nt database were used as input for blobtools v1.1.1 (Laetsch and Blaxter 2017) and scaffolds identified as possible contamination were removed from the assembly. Finally, purge_dups (Guan et al. 2020) v1.2.5 was used to remove haplotigs and contig overlaps. The primary assembly was scaffolded by Dovetail using the OmniC reads with the HiRise assembler (Putnam et al. 2016), after aligning the OmniC library reads to the filtered draft input assembly using bwa v0.7.15. (Manni et al. 2021) using both the *Embryophyta* and *Eukaryota* databases.

The separations of Dovetail OmniC read pairs mapped within draft scaffolds were analyzed by HiRise to produce a likelihood model for genomic distance between read pairs, and the model was used to identify and break putative misjoins, to score prospective joins, and make joins above a threshold.

Assembly completeness was assessed using BUSCO v. 5.4.4 (Manni et al. 2021).

### Sex-linked SNP identification

We mapped RNAseq leaf expression data from population samples of TX and NC cytotype male and female plants of *R. hastatulus* (Hough et al. 2014) to both haplotype assemblies using STAR v2.7.10a (Dobin et al. 2013). We performed variant calling using freebayes v1.3.4 (Garrison and Marth 2012), then filtered sites to a final set comprised of biallelic sites with genotype quality > 30. We then used custom R scripts to identify putative sex-linked SNPs. We selected all sites that were heterozygous in all 6 males and homozygous in all 6 females per cytotype to obtain candidate fixed SNP differences between X and Y.

### Gene annotation

Gene annotation followed previous approaches (Rifkin et al. 2022). In particular, we performed the annotation with MAKER-3.01.03 (Cantarel et al. 2008) in four rounds. In the first round, we used the *Rumex* RNA-Seq transcripts from previously published floral and leaf transcriptomes (Hough et al. 2014; Sandler et al. 2018) and annotated Tartary buckwheat proteins from version FtChromosomeV2.IGDBv2 (Zhang et al. 2017) for inferring gene predictions; and used the transposable element (TE) library (see below) to mask the genome. We trained the resulting annotation for SNAP gene predictor, using the gene models with an AED of 0.5 or better and a length of 50 or more amino acids. In the following rounds, we used the resulting EST and protein alignments from the first round, and the SNAP model from the previous round for annotation. We functionally annotated the final gene models based on BLAST v 2.2.28+ (Altschul et al. 1990) and InterProScan 5.52–86.0 (Jones et al. 2014), by using the related scripts in the Maker package.

### Syntenic gene alignments and analysis

We estimated orthology and synteny between protein coding genes in haplotype 1, haplotype 2 and *R. salicifolius* using the R package GENESPACEv1.1.8 (Lovell et al. 2022), which uses MCScanX (Wang et al. 2012) to infer syntenic gene blocks and then implements ORTHOFINDERv2.5.4 (Emms and Kelly 2019) and DIAMONDv2.1.4.158 (Buchfink et al. 2021) to find orthogroups within syntenic blocks. We performed analyses and visualized results in Rv4.1.0 (R Core Team et al. 2022). We used default parameters, with the exception of ORTHOFINDER one-way sequence search, which is appropriate for our closely related genomes.

We also conducted whole-genome pairwise alignments between the two haplotypes using Anchorwave v. 1.01 (Song et al. 2022), using the options allowing for relocation variation, and chromosome fusion. We used Minimap2 in the Anchorwave alignment, followed by Proalign using “-Q 1” option.

### Ks analysis

We calculated synonymous substitution rate between paternal and maternal haplotype assemblies using SynMap2 on the COGE platform (Haug-Baltzell et al. 2017). To compare homologous genes between haplotypes we used a cut-off of K<0.2. We plotted median *Ks* values in 100 gene sliding widows (step size =1) relative to their positions on the X chromosome.

### Gene gain and loss

Pangenome annotations produced by GENESPACE provide a list of orthologous genes shared by each genome, and their positions relative to an assigned reference genome. We excluded genes with non-syntenic orthologs, and genes belonging to arrays that were not defined as representative by GENESPACE from subsequent analysis. We calculated the number of genes lost on the paternal or maternal assembly by counting the number of syntenic genes found in both *R. salicifolius* and the other phased haplotype but absent from the focal haplotype assembly. To determine whether candidate lost genes are indeed lost and not simply missing from the annotation, we performed BLASTv2.5.0+ (Altschul et al. 1990) of these gene transcripts to the entire genome assembly sequence. We selected only the top BLAST hit (by percent identity) per candidate lost gene. We classified genes as present if the top BLAST sequence was on the corresponding chromosome. We classified genes not meeting this condition as lost, as well as genes where less than 50% of the query sequence is aligned to the subject. We defined these genes with less than 50% of query aligned as partially lost and included them within the total number of lost genes.

### Non-syntenic orthologs

We identified one-to-one orthologs within the pangenome annotation where a syntenic ortholog was shared with the outgroup, *R. salicifolius*, in only one of the haplotypes, while the other haplotype’s ortholog was non-syntenic, as defined by GENESPACE. To determine whether there is an association between sex-linked regions and haplotype in terms of non-syntenic ortholog content, we performed a 2×2 chi-square test of independence (R v4.3.1) comparing counts in the old sex-linked region and all autosomes for both haplotypes.

### Satellite identification, TE annotation and analysis

We identified satellites using RepeatExplorer2 (Novák et al. 2020). We then pre-processed short-read Illumina data (Beaudry et al. 2017) with RepeatExplorer’s inbuilt preprocessing pipeline (Novák et al. 2013, 2020). Trimming step was set to keep only full length 150 bp reads and discard low quality reads (quality cutoff=10, percent above cutoff=95) or reads containing adapters. We ran RepeatExplorer2/TAREAN pipeline (version 0.3.8-451, Novák at al. 2017, 2020) with default parameters. To compare repeats in the 4 samples, a comparative analysis was run following Novák et al. (2020), where we analyzed reads from all samples together with equal coverage between samples (using genome sizes according to Grabowska-Joachimiak et al. 2015). We further used sex and cytotype specific clusters for cytogenetic analysis (Table S7).

We produced the transposable element annotation using EDTA (Extensive de-novo TE Annotator) v. 2.1.0 pipeline (Ou et al. 2019). This pipeline combines the best-performing structure- and homology-based TE finding programs (GenomeTools, LTR_FINDER_parallel (Ou and Jiang 2019), LTR_harvest_parallel, LTR_retriever (Ou and Jiang 2018), Generic Repeat Finder (Shi and Liang 2019), TIR-Learner (Su et al. 2019), HelitronScanner (Xiong et al. 2014), TEsorter (Zhang et al. 2022) and filters their results to produce a comprehensive and non-redundant TE library. We used the optional parameters ‘–sensitive 1’ and ‘–anno 1’ to identify remaining unidentified TEs with RepeatModeler and to produce an annotation. We used custom R scripts to visualise the data.

To analyse insertions near genes, we used Bedtools v2.30.0 and custom R scripts to compare the TE annotation file against the gene annotation, using genes lifted over from the paternal haplotype to the maternal haplotype with Liftoff 1.6.3 (Shumate and Salzberg 2020).

### Chromosome preparation and cytogenetic analysis

We used young seedlings of *R. hastatulus* of both XY, XYY cytotypes (North Carolina and Texas) for chromosome preparation, cell synchronization and metaphase chromosome arrest as described in Bačovský et al. (2020). Additionally, we grew plants of XYY cytotype in a hydroponic tank with Hoagland solution in a growth chamber with a 16 h light/8 h dark cycle at 22°C (Hoagland & Snyder, 1933). We collected young roots from hydroponic tanks once per every two weeks, synchronised in ice-cold water at 4°C for 24 to 28 h. After the cell synchronization, we immediately fixed the root tissue was in Clarke’s fixative (ethanol:glacial acetic acid, 3:1, v:v) and stored it at 37°C. After one week of fixation, we replaced the fixative, and fixed roots were stored at −20°C until further use.

We isolated the DNA used for PCR amplification from young leaves using CTAB (cetyltrimethylammonium bromide) solution and chloroform. We ground young leaves in a sterile grinder in liquid nitrogen and added 1 ml of CTAB solution to each sample. We vortexed the mixture for 30 s and incubated it at 65 °C for 45 min. We then added 2 μl of RnaseA (concentration 200 mg/ml) for the last 5 min. Next, we added 700 μl of chloroform and acetic acid solution (chloroform:glacial acetic acid, 24:1, v:v) to each sample, vortexed each for 1 min and centrifuged at maximum speed (14 000 rpm) for 2 min. We transferred the upper aqueous layer to a new tube and added 700 μl of chloroform, then vortexed and centrifuged the samples for an additional 5 min at 14 000 rpm. We again transferred the upper aqueous layer to a new tube, and precipitated resulting DNA with 800 μl of isopropanol. We vortexed and centrifuged the mixture at maximum speed for additional 5 min, then discarded the supernatant, and repeated the whole step using 75% ethanol to remove any excess salts. Finally, we air-dried the pellet for 5 min and dissolved in 20-40 μl of 1X Tris EDTA buffer for 45 minutes. We then analyzed isolated DNA on 1% agarose gel, and its concentration and purity were measured on Nanodrop.

Based on the RepeatExplorer analysis (Figure S2), we designed new primers for XY/XYY cytotype and sex-specific satellites in GeneiousPrime (2023.1.1) and synthesized in GeneriBiotech. We used primers directly for PCR amplification (Table S7), and we amplified the satellites to the manufacturer’s instructions using 0.4 μl of *R. hastatulus* gDNA (TopBio, Vestec, Czech Republic, T034). The PCR conditions were as follows: 4 min at 94°C, 36 cycles of 20 s at 94°C, 20 s at 50 – 60°C (Table S7), 30 s at 72°C (for Cl135 1 min) and final extension 5 min at 72°C. We verified the PCR products by 1% agarose electrophoresis with EtBr staining and purified using the QiaQuick PCR Purification Kit (Qiagen, Hilden, Germany, 28104) following manufacturer’s instructions. We then verified the purified products again by agarose electrophoresis, followed by Nanodrop measurement.

We labelled the purified DNA by nick translation according to the manufacturer’s instructions using Atto488 NT (PP-305L-488), Atto550 NT (PP-305L-550) or Cy5 (PP-305L-647N) (Jena Bioscience, Jena, Germany). The reaction proceeded for 1 hour and 30 minutes at 15°C. We verified the nicked-DNA products on 1% agarose gel with EtBr staining. We then placed the reactions on ice directly after the reaction to avoid the over-labelling of DNA before addition of EDTA. We used the nick translated products as DNA probes in FISH (fluorescence in situ hybridisation). The hybridisation mixture (87 % stringency; Table S8) included 1 μl of labelled DNA (the final volume 1.5 ng/μl). We carefully mixed the hybridisation mixture and denatured the sample at 85°C for 10 minutes, transferred on ice for 5 minutes to perform FISH.

We prepared chromosomes using the squashing technique as described in Karafiátová et al. (2016) and Bačovský et al. (2020) with minor modifications, using 0.05M HCl acid in 0.001 M citrate buffer before enzymatic digestion. We used slides containing chromosomes with well-preserved morphology and structure for FISH, as described in Schubert et al. (2016) with minor modifications. Briefly, we first washed the slides for 2× 5 min in 2× SSC solution (pH 7.2 – 7.5) and then refixed the slides in Clarke’s fixative for 10 min, and washed 2× 5 min in 2× SSC solution. To remove the remnants of cytoplasm, we treated slides with pepsin (50 mg/ml) diluted in 2× SSC in a water bath at 37°C for 5 to 15 min. Next, we washed the slides for 2× 5 min in 2× SSC solution and re-fixed them for 10 min in 3.7% formaldehyde (diluted in 2× SSC). After fixation, we washed the slides for 2× 5 min in 2× SSC solution, shortly washed in distilled water and dehydrated in an ethanol series (60%, 80%, 100%), each step 2 min. We then applied denatured hybridisation mixture (20 ml each) to each slide, covered with coverslip, placed on a hot plate at 77°C for 2 min and transferred at 37°C overnight. We washed slides for 5 min in 2× SSC, transferred for 20 min at 57°C in 2× SSC, washed 5 min in 2× SSC at RT again, and dehydrated in an ethanol series (60%, 80%, 100%). Finally, we mounted the slides in VectaShield (Vector, H-1500) supplemented with DAPI (2-(4-aminophenyl)-1H-indole-6-carboxamidine). We captured chromosomes under an Olympus AX70 fluorescence microscope equipped with a CCD camera, and Imaris software. We used the software GIMP-2.10 and Affinity Photo 2 to process all channels.

## Data Availability

Final assemblies are uploaded onto COGE (https://genomevolution.org/coge/) and will be accessible at xxxx. All raw reads will be uploaded to the SRA under accession number xxxx. All custom R Scripts are available on Github (https://github.com/SIWLab/XYYmaleGenome).

## Acknowledgements

We thank Meng Yuan, Bill Cole, and Thomas Gludovacz for help with plant growth and maintenance, and Alex Harkess and Sarah Carey for discussion and advice on sex chromosome assembly methods. This research was funded by Discovery Grants from the Natural Sciences and Engineering Research Council of Canada to SCHB and SIW.

## Notes

### Competing Interest Statement

The authors have declared no competing interest.

### Summary of Updates

Added cytogenetic results, additional discussion, and corrected wording.

## References

Abbott JK, Nordén AK, Hansson B. 2017. Sex chromosome evolution: historical insights and future perspectives. Proc. Biol. Sci. 284: 20162806

Akagi T, Fujita N, Masuda K, Shirasawa K, Nagaki K, Horiuchi A, Kuwada E, Kunou R, Nakamura K, Ikeda Y, et al. 2023. Rapid and dynamic evolution of a giant Y chromosome in *Silene latifolia*. bioRxiv [Internet]:2023.09.21.558759.

Altschul SF, Gish W, Miller W, Myers EW, Lipman DJ. 1990. Basic local alignment search tool. J. Mol. Biol. 215:403–410.

Bachtrog D. 2013. Y-chromosome evolution: emerging insights into processes of Y-chromosome degeneration. Nat. Rev. Genet. 14:113–124.

Bachtrog D. 2020. The Y Chromosome as a battleground for intragenomic conflict. Trends Genet. 36:510–522.

Bačovský V, Čegan R, Šimoníková D, Hřibová E, & Hobza R. 2020. The formation of sex chromosomes in *Silene latifolia* and *S. dioica* was accompanied by multiple chromosomal rearrangements. Front. Plant Sci. 11:205

Beaudry FEG, Barrett SCH, Wright SI. 2017. Genomic loss and silencing on the Y chromosomes of *Rumex*. Genome Biol. Evol. 9:3345–3355.

Beaudry FEG, Barrett SCH, Wright SI. 2020. Ancestral and neo-sex chromosomes contribute to population divergence in a dioecious plant. Evolution 74:256–269.

Beaudry FEG, Rifkin JL, Peake AL, Kim D, Jarvis-Cross M, Barrett SCH, Wright SI. 2022. Effects of the neo-X chromosome on genomic signatures of hybridization in *Rumex hastatulus*. Mol. Ecol. 31:3708–3721.

Bergero R, Qiu S, Charlesworth D. 2015. Gene loss from a plant sex chromosome system. Curr. Biol. 25:1234–1240.

Bieker VC, Battlay P, Petersen B, Sun X, Wilson J, Brealey JC, Bretagnolle F, Nurkowski K, Lee C, Barreiro FS, et al. 2022. Uncovering the genomic basis of an extraordinary plant invasion. Sci Adv 8:eabo5115.

Buchfink B, Reuter K, Drost H-G. 2021. Sensitive protein alignments at tree-of-life scale using DIAMOND. Nat. Methods 18:366–368.

Cantarel BL, Korf I, Robb SMC, Parra G, Ross E, Moore B, Holt C, Sánchez Alvarado A, Yandell M. 2008. MAKER: an easy-to-use annotation pipeline designed for emerging model organism genomes. Genome Res. 18:188–196.

Charlesworth D, Charlesworth B, Marais G. 2005. Steps in the evolution of heteromorphic sex chromosomes. Heredity 95:118–128.

Cheng H, Concepcion GT, Feng X, Zhang H, Li H. 2021. Haplotype-resolved de novo assembly using phased assembly graphs with hifiasm. Nat. Methods 18:170–175.

Crowson D, Barrett SCH, Wright SI. 2017. Purifying and positive selection influence patterns of gene loss and gene expression in the evolution of a plant sex chromosome system. Mol. Biol. Evol. 34:1140–1154.

Cuñado N, Navajas-Pérez R, de la Herrán R. et al. 2007. The evolution of sex chromosomes in the genus *Rumex* (Polygonaceae): Identification of a new species with heteromorphic sex chromosomes. Chromosome Res. 15: 825–833.

Dobin A, Davis CA, Schlesinger F, Drenkow J, Zaleski C, Jha S, Batut P, Chaisson M, Gingeras TR. 2013. STAR: ultrafast universal RNA-seq aligner. Bioinformatics 29:15–21.

Durand NC, Robinson JT, Shamim MS, Machol I, Mesirov JP, Lander ES, Aiden EL. 2016. Juicebox provides a visualization system for Hi-C contact maps with unlimited zoom. Cell Syst 3:99–101.

Ellinghaus D, Kurtz S, Willhoeft U. 2008. LTRharvest, an efficient and flexible software for de novo detection of LTR retrotransposons. BMC Bioinformatics 9:18.

Emms DM, Kelly S. 2019. OrthoFinder: phylogenetic orthology inference for comparative genomics. Genome Biol. 20:238.

Fuller ZL, Koury SA, Phadnis N, Schaeffer SW. 2019. How chromosomal rearrangements shape adaptation and speciation: Case studies in *Drosophila pseudoobscura* and its sibling species *Drosophila persimilis*. Mol. Ecol. 28:1283–1301.

Garrison E, Marth G. 2012. Haplotype-based variant detection from short-read sequencing. arXiv [q-bio.GN] [Internet]. Available from: http://arxiv.org/abs/1207.3907

Grabowska-Joachimiak A, Kula A, Książczyk T, Chojnicka J, Sliwinska E, Joachimiak AJ. 2015. Chromosome landmarks and autosome-sex chromosome translocations in *Rumex* hastatulus, a plant with XX/XY1Y2 sex chromosome system. Chromosome Res. 23:187–197.

Guan D, McCarthy SA, Wood J, Howe K, Wang Y, Durbin R. 2020. Identifying and removing haplotypic duplication in primary genome assemblies. Bioinformatics 36:2896–2898.

Haug-Baltzell A, Stephens SA, Davey S, Scheidegger CE, Lyons E. 2017. SynMap2 and SynMap3D: web-based whole-genome synteny browsers. Bioinformatics 33:2197– 2198.

Hoagland DR, Snyder WC. 1933. Nutrition of strawberry plant under controlled conditions. (a) Effects of deficiencies of boron and certain other elements, (b) Susceptibility to injury from sodium salts. J. Am. Soc. Hortic. Sci. 30: 288–294.

Hough J, Hollister JD, Wang W, Barrett SCH, Wright SI. 2014. Genetic degeneration of old and young Y chromosomes in the flowering plant *Rumex hastatulus*. Proc. Natl. Acad. Sci. U. S. A. 111:7713–7718.

Jones P, Binns D, Chang H-Y, Fraser M, Li W, McAnulla C, McWilliam H, Maslen J, Mitchell A, Nuka G, et al. 2014. InterProScan 5: genome-scale protein function classification. Bioinformatics 30:1236–1240.

Karafiátová M, Bartoš J, & Doležel J. 2016. Localization of low-copy DNA sequences on mitotic chromosomes by FISH. Methods Mol Biol 1429:49–64.

Kasjaniuk M, Grabowska-Joachimiak A, Joachimiak AJ. 2019. Testing the translocation hypothesis and Haldane’s rule in *Rumex hastatulus*. Protoplasma 256:237–247.

Kent TV, Uzunović J, Wright SI. 2017. Coevolution between transposable elements and recombination. Philos. Trans. R. Soc. Lond. B Biol. Sci. 372: 20160458

Kirkpatrick M, Barton N. 2006. Chromosome inversions, local adaptation and speciation. Genetics 173:419–434.

Laetsch DR, Blaxter ML. 2017. BlobTools: Interrogation of genome assemblies. F1000Res. 6:1287.

Lenormand T, Fyon F, Sun E, Roze D. 2020. Sex chromosome degeneration by regulatory evolution. Curr. Biol. 30 (15):3001–3006

Li H. 2018. Minimap2: pairwise alignment for nucleotide sequences. Bioinformatics 34:3094–3100.

Li H, Durbin R. 2009. Fast and accurate short read alignment with Burrows-Wheeler transform. Bioinformatics 25:1754–1760.

Löve Á. 1986. Chromosome Number Reports XCII. Taxon 35:610–613.

Lovell JT, Sreedasyam A, Schranz ME, Wilson M, Carlson JW, Harkess A, Emms D, Goodstein DM, Schmutz J. 2022. GENESPACE tracks regions of interest and gene copy number variation across multiple genomes. Elife 11:e78526.

Lowry DB, Willis JH. 2010. A widespread chromosomal inversion polymorphism contributes to a major life-history transition, local adaptation, and reproductive isolation. PLoS Biol. 8(9): e1000500

Mandáková T, Pouch M, Brock JR, Al-Shehbaz IA, Lysak MA. 2019. Origin and evolution of diploid and allopolyploid *Camelina* genomes were accompanied by chromosome shattering. Plant Cell 31:2596–2612.

Manni M, Berkeley MR, Seppey M, Simão FA, Zdobnov EM. 2021. BUSCO update: novel and streamlined workflows along with broader and deeper phylogenetic coverage for scoring of eukaryotic, rrokaryotic, and viral genomes. Mol. Biol. Evol. 38:4647–4654.

Mrnjavac A, Khudiakova KA, Barton NH, Vicoso B. 2023.Slower-X: reduced efficiency of selection in the early stages of X chromosome evolution. Evol Lett 7:4–12.

Muyle AM, Seymour DK, Lv Y, Huettel B, Gaut BS. 2022. Gene body methylation in plants: mechanisms, functions, and important implications for understanding evolutionary processes. Genome Biol. Evol. 14(4):evac038

Novák P, Neumann P, Pech J, Steinhaisl J, Macas J. 2013. RepeatExplorer: a Galaxy-based web server for genome-wide characterization of eukaryotic repetitive elements from next-generation sequence reads. Bioinformatics 29:792–793.

Novák P, Ávila Robledillo L, Koblížková A, Vrbová I, Neumann P, Macas J. 2017. TAREAN: a computational tool for identification and characterization of satellite DNA from unassembled short reads. Nucleic Acids Res. 45:e111.

Novák P, Neumann P, Macas J. 2020. Global analysis of repetitive DNA from unassembled sequence reads using RepeatExplorer2. Nat. Protoc. 15:3745–3776.

Orr HA, Kim Y. 1998. An adaptive hypothesis for the evolution of the Y chromosome. Genetics 150:1693–1698.

Ou S, Jiang N. 2018. LTR_retriever: a highly accurate and sensitive program for identification of long terminal repeat retrotransposons. Plant Physiology 176:1410– 1422.

Ou S, Jiang N. 2019. LTR_FINDER_parallel: parallelization of LTR_FINDER enabling rapid identification of long terminal repeat retrotransposons. Mob. DNA 10:48.

Ou S, Su W, Liao Y, Chougule K, Agda JRA, Hellinga AJ, Lugo CSB, Elliott TA, Ware D, Peterson T, et al. 2019. Benchmarking transposable element annotation methods for creation of a streamlined, comprehensive pipeline. Genome Biol. 20:275.

Papadopulos AST, Chester M, Ridout K, Filatov DA. 2015. Rapid Y degeneration and dosage compensation in plant sex chromosomes. Proc. Natl. Acad. Sci. U. S. A. 112:13021–13026.

Peichel CL, McCann SR, Ross JA, Naftaly AFS, Urton JR, Cech JN, Grimwood J, Schmutz J, Myers RM, Kingsley DM, et al. 2020. Assembly of the threespine stickleback Y chromosome reveals convergent signatures of sex chromosome evolution. Genome Biol. 21:177.

Pickup M, Barrett SCH. 2013. The influence of demography and local mating environment on sex ratios in a wind-pollinated dioecious plant. Ecol. Evol. 3:629–639.

Putnam NH, O’Connell BL, Stites JC, Rice BJ, Blanchette M, Calef R, Troll CJ, Fields A, Hartley PD, Sugnet CW, et al. 2016. Chromosome-scale shotgun assembly using an in vitro method for long-range linkage. Genome Res. 26:342–350.

R Core Team R, Others. 2022.R: A language and environment for statistical computing.

Rice WR. 1987. The accumulation of sexually antagonistic genes as a selective agent promoting the evolution of reduced recombination between primitive sex chromosomes. Evolution 41:911–914.

Rifkin JL, Beaudry FEG, Humphries Z, Choudhury BI, Barrett SCH, Wright SI. 2021. Widespread recombination suppression facilitates plant sex chromosome evolution. Mol. Biol. Evol. 38:1018–1030.

Rifkin JL, Hnatovska S, Yuan M, Sacchi BM, Choudhury BI, Gong Y, Rastas P, Barrett SCH, Wright SI. 2022. Recombination landscape dimorphism and sex chromosome evolution in the dioecious plant *Rumex hastatulus*. Philos. Trans. R. Soc. Lond. B Biol. Sci. 377:20210226.

Sandler G, Beaudry FEG, Barrett SCH, Wright SI. 2018. The effects of haploid selection on Y chromosome evolution in two closely related dioecious plants. Evol Lett 2:368– 377.

Schubert V, Ruban A, & Houben A. 2016. Chromatin ring formation at plant centromeres. Front. Plant Sci. 7:28.

Shi J, Liang C. 2019. Generic Repeat Finder: a high-sensitivity tool for genome-wide *de novo* repeat detection. Plant Physiol. 180:1803–1815.

Shumate A, Salzberg SL. 2020. Liftoff: an accurate gene annotation mapping tool. bioRxiv 2020.06.24.169680. Available from: https://www.biorxiv.org/content/10.1101/2020.06.24.169680

Smith BW. 1964. The evolving karyotype of *Rumex hastatulus*. Evolution 18:93–104.

Song B, Marco-Sola S, Moreto M, Johnson L, Buckler ES, Stitzer MC. 2022. AnchorWave: Sensitive alignment of genomes with high sequence diversity, extensive structural polymorphism, and whole-genome duplication. Proc. Natl. Acad. Sci. U. S. A. 119 (1) e2113075119

Su W, Gu X, Peterson T. 2019. TIR-Learner, a new ensemble method for TIR transposable element annotation, provides evidence for abundant new transposable elements in the maize genome. Mol. Plant 12:447–460.

Subrini J, Turner J. 2021. Y chromosome functions in mammalian spermatogenesis. Elife 10:e67345.

Todesco M, Owens GL, Bercovich N, Légaré J-S, Soudi S, Burge DO, Huang K, Ostevik KL, Drummond EBM, Imerovski I, et al. 2020. Massive haplotypes underlie ecotypic differentiation in sunflowers. Nature 584:602–607.

Wang Y, Tang H, Debarry JD, Tan X, Li J, Wang X, Lee T-H, Jin H, Marler B, Guo H, et al. 2012. MCScanX: a toolkit for detection and evolutionary analysis of gene synteny and collinearity. Nucleic Acids Res. 40:e49.

Wei KH-C, Gibilisco L, Bachtrog D. 2020. Epigenetic conflict on a degenerating Y chromosome increases mutational burden in Drosophila males. Nat. Commun. 11:5537.

Wicker T, Sabot F, Hua-Van A, Bennetzen JL, Capy P, Chalhoub B, Flavell A, Leroy P, Morgante M, Panaud O, et al. 2007. A unified classification system for eukaryotic transposable elements. Nat. Rev. Genet. 8:973–982.

Xiong W, He L, Lai J, Dooner HK, Du C. 2014. HelitronScanner uncovers a large overlooked cache of Helitron transposons in many plant genomes. Proceedings of the National Academy of Sciences 111:10263–10268.

Zhang L, Li X, Ma B, Gao Q, Du H, Han Y, Li Y, Cao Y, Qi M, Zhu Y, et al. 2017. The Tartary buckwheat genome provides insights into rutin biosynthesis and abiotic stress tolerance. Mol. Plant 10:1224–1237.

Zhang R-G, Li G-Y, Wang X-L, Dainat J, Wang Z-X, Ou S, Ma Y. 2022. TEsorter: an accurate and fast method to classify LTR-retrotransposons in plant genomes. Hortic Res 9: uhac017.

Zhou C, McCarthy SA, Durbin R. 2023. YaHS: yet another Hi-C scaffolding tool. Bioinformatics 39(1): btac808

