## Supplementary Information for "Phased assembly of neo-sex chromosomes reveals extensive Y degeneration and rapid genome evolution in *Rumex hastatulus*"

#### Table S1- Genome Assembly Statistics

| Assembly | Total Assembly Length | Contig N50 | Contig L50 | Scaffold N50 | Scaffold L50 | Scaffold N90 | Scaffold L90 |
|---|---|---|---|---|---|---|---|
| *R. salicifolius* | 586,431,610 | 53,106,915 | 5 | 56,809,946 | 5 | 51,010,208 | 9 |
| *R. hastatulus* maternal haplotype | 1,510,206,782 | 6,283,184 | 73 | 466,657,150 | 2 | 168,693,024 | 4 |
| *R. hastatulus* paternal haplotype | 1,719,830,090 | 5,620,551 | 92 | 343,398,141 | 3 | 166,046,2686 | 5 |

#### Table S2 - BUSCO results

| **Genome** | **OrthoDB** | **Complete, single copy** | **Complete, duplicated** | **Fragmented** | **Missing** |
|---|---|---|---|---|---|
| *R. salicifolius* | Eukaryota | 227 (89%) | 27 (10.6%) | 0 | 1 (0.4%) |
| *R. hastatulus* maternal haplotype | Eukaryota | 223 (87.5%) | 30 (11.8%) | 0 | 2 (0.7%) |
| *R. hastatulus* paternal haplotype | Eukaryota | 224 (87.8%) | 30 (11.8%) | 0 | 1 (0.4%) |
| *R. salicifolius* | Embryophyta | 1528 (94.7%) | 39 (2.4%) | 20 (1.2%) | 27 (1.7%) |
| *R. hastatulus* maternal haplotype | Embryophyta | 1476 (91.4%) | 77 (4.8%) | 20 (1.2%) | 41 (2.6%) |
| *R. hastatulus* paternal haplotype | Embryophyta | 1463 (90.6%) | 71 (4.4%) | 26 (1.6%) | 54 (3.4%) |

### Table S3 – Non-syntenic orthologs

| Region | Number of non-syntenic orthologs unique to maternal haplotype | Number of non-syntenic orthologs unique to paternal haplotype |
|---|---|---|
| A1 | 14 | 22 |
| A2 | 11 | 34 |
| A4 | 6 | 6 |
| New sex-linked region | 1 | 6 |
| Old sex-linked region | 13 | 155 |
| Pseudo-autosomal regions | 2 | 15 |

### Table S4 - TE count chi-square

| Region | Linkage | p-value | Expected | X observed | Y observed | Y/X ratio |
|---|---|---|---|---|---|---|
| entire | New | 1.332333e-71 | 88431.5 | 92194 | 84669 | 0.9183786 |
| entire | Old | 0.00000e+00 | 299641.5 | 158443 | 440840 | 2.7823255 |
| entire | PAR | 2.220696e-11 | 160351.5 | 158457 | 162246 | 1.0239118 |
| upstream | New | 8.139606e-02 | 951.0 | 913 | 989 | 1.0832421 |
| upstream | Old | 5.521570e-27 | 1608.0 | 1303 | 1913 | 1.4681504 |
| upstream | PAR | 1.598126e-05 | 5035.5 | 4819 | 5252 | 1.0898527 |
| downstream | New | 5.394717e-06 | 1015.5 | 913 | 1118 | 1.2245345 |
| downstream | Old | 2.074799e-41 | 1774.5 | 1373 | 2176 | 1.5848507 |
| downstream | PAR | 2.162392e-16 | 5045.5 | 4633 | 5458 | 1.1780704 |
| exon | New | 3.411261e-01 | 565.0 | 549 | 581 | 1.0582878 |
| exon | Old | 5.152438e-19 | 1154.0 | 940 | 1368 | 1.4553191 |
| exon | PAR | 2.187631e-20 | 1703.0 | 1973 | 1433 | 0.7263051 |
| intron | New | 2.398108e-02 | 812.5 | 767 | 858 | 1.1186441 |
| intron | Old | 1.104104e-19 | 1334.5 | 1100 | 1569 | 1.4263636 |
| intron | PAR | 6.717053e-01 | 4454.0 | 4474 | 4434 | 0.9910595 |

#### Table S5 – XYY male paternal haplotype gene loss results

| Region | Number of genes with syntenic orthologs in outgroup | Proportion of genes completely absent in paternal haplotype | Proportion of genes partially absent in paternal haplotype |
|---|---|---|---|
| A1 | 2786 | 0.0004 | 0.0262 |
| A2 | 2833 | 0.0039 | 0.0289 |
| A4 | 1535 | 0.0033 | 0.0319 |
| New X | 252 | 0.0000 | 0.0278 |
| Old X | 745 | 0.2107 | 0.1275 |
| PAR1 | 793 | 0.0076 | 0.0303 |
| PAR2 | 1295 | 0.0015 | 0.0409 |

#### Table S6 – XYY male maternal haplotype gene loss results

| Region | Number of genes with syntenic orthologs in outgroup | Proportion of genes absent in maternal haplotype |
|---|---:|---:|
| A1 | 2702 | 0.0248 |
| A2 | 2714 | 0.0247 |
| A4 | 1451 | 0.0207 |
| NewY1 | 145 | 0.0069 |
| NewY2 | 96 | 0.0208 |
| OldY1 | 206 | 0.0146 |
| OldY2 | 280 | 0.0250 |
| PAR1 | 755 | 0.0265 |
| PAR2 | 1236 | 0.0218 |

#### Table S7 The list of used primers of candidate satellites

| Primer | Sequence (5´-3´) | Product size | Annealing temperature (°C) |
|---|---|---|---|
| **Cl12F** | AGTTCGTTGTCCAATATACTCGTT | 117 | 58 |
| **Cl12R** | TGTGAAAACGTACCGAACATGT | | |
| **Cl133 F** | GTATAAAAACCGTAACCGTG | 66 | 53 |
| **Cl133 R** | GCTCGAAATAGTGTAATCAAAC | | |
| **Cl134 5S F** | CCACTGATATATTGACCGCTCGA | 265 | 60 |
| **Cl134 5S R** | AAAGGCAATATCTCATCCGATAGT | | |
| **Cl135 F** | CAATTTCATGACGGTTAATAACG | 1136 | 55 |
| **Cl135 R** | CACGGTTACGGTTTTTATAC | | |
| **Cl162F** | GCTCAAAATTTTGTTGCTCTCC | 80 | 60 |
| **Cl162R** | TGGTGACTTAAAACATCAGAGCAAC | | |
| **Cl168 F** | TGTACAATGGTAGAAATCGGGTC | 170 | 55 |
| **Cl168 R** | CAAATTTTGTTCAATCTC | | |
| **Cl86 F** | CACTTGCCCGATGGAAACGGC | 170 | 55 |
| **Cl86 R** | CCTTGTTCATCCGTTTCCGGTG | | |

Table S8 The composition of the hybridization mixture used in FISH

| Hybridization mixture Stringency 87% | Volume per one slide (µl) |
|---|---|
| **100% formamide** | 10 |
| **50% dextran sulphate** | 4 |
| **20xSSC** | 0.5 |
| **ddH$_2$O** | X (to final volume) |
| **DNA probe** | 1 (for each cluster) |
| **Final volume 20 µl** | |

**Figure S1**. Sex-linked SNP patterns along the XYY cytotype's X, Y1 and Y2 chromosomes. Density of sex-linked SNPs in 1 MB windows in the XY and XYY cytotypes. Dashed line indicates the inferred breakpoint between the old X (Y) chromosome and the neo-X (neo-Y) region.

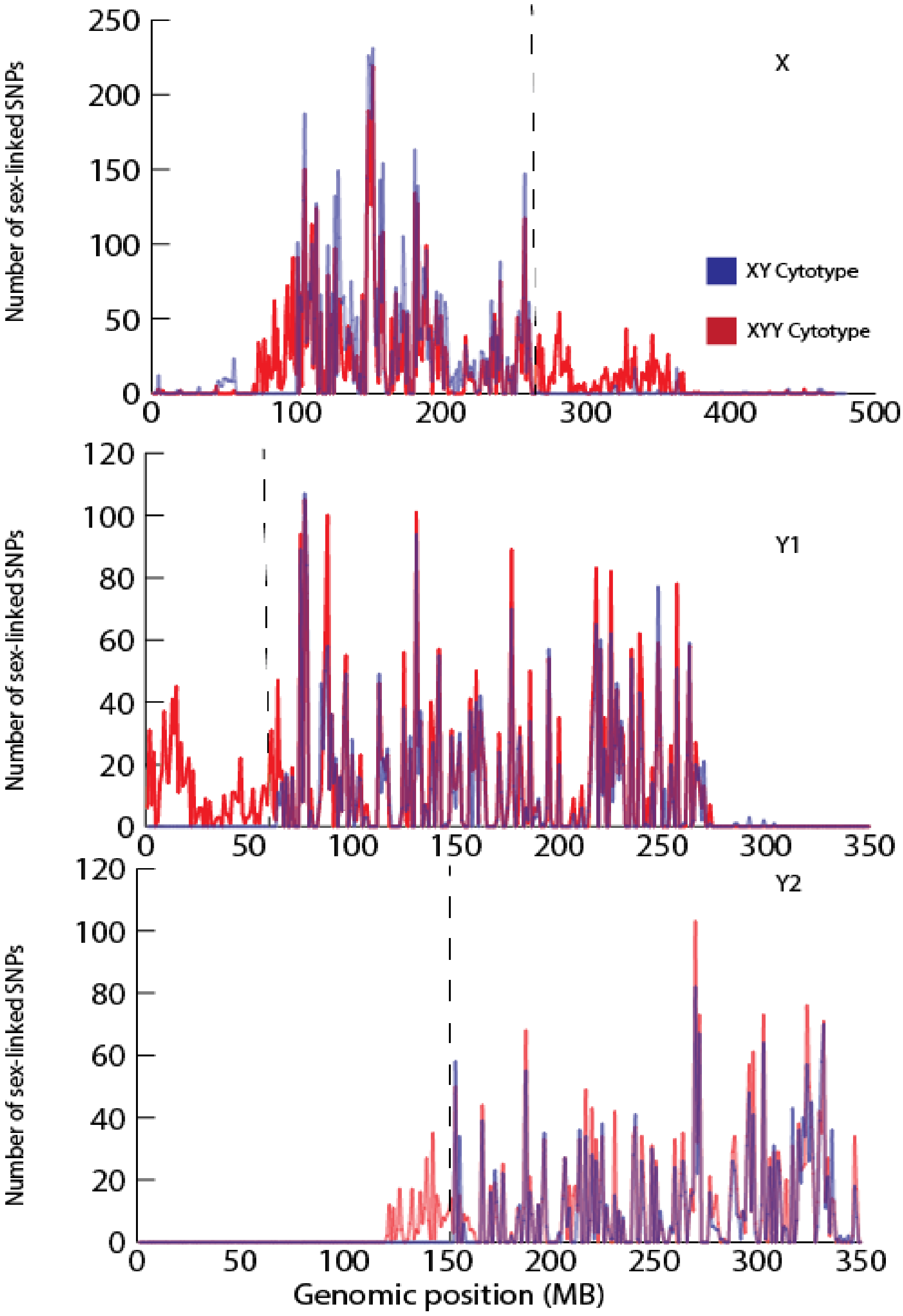

**Figure S2**. Repeat cluster abundance in XYY and XY cytotype. Comparison of males and female (top) and XYY/XY cytotypes (bottom). Right plots are identical to left plots in logarithmic scale.

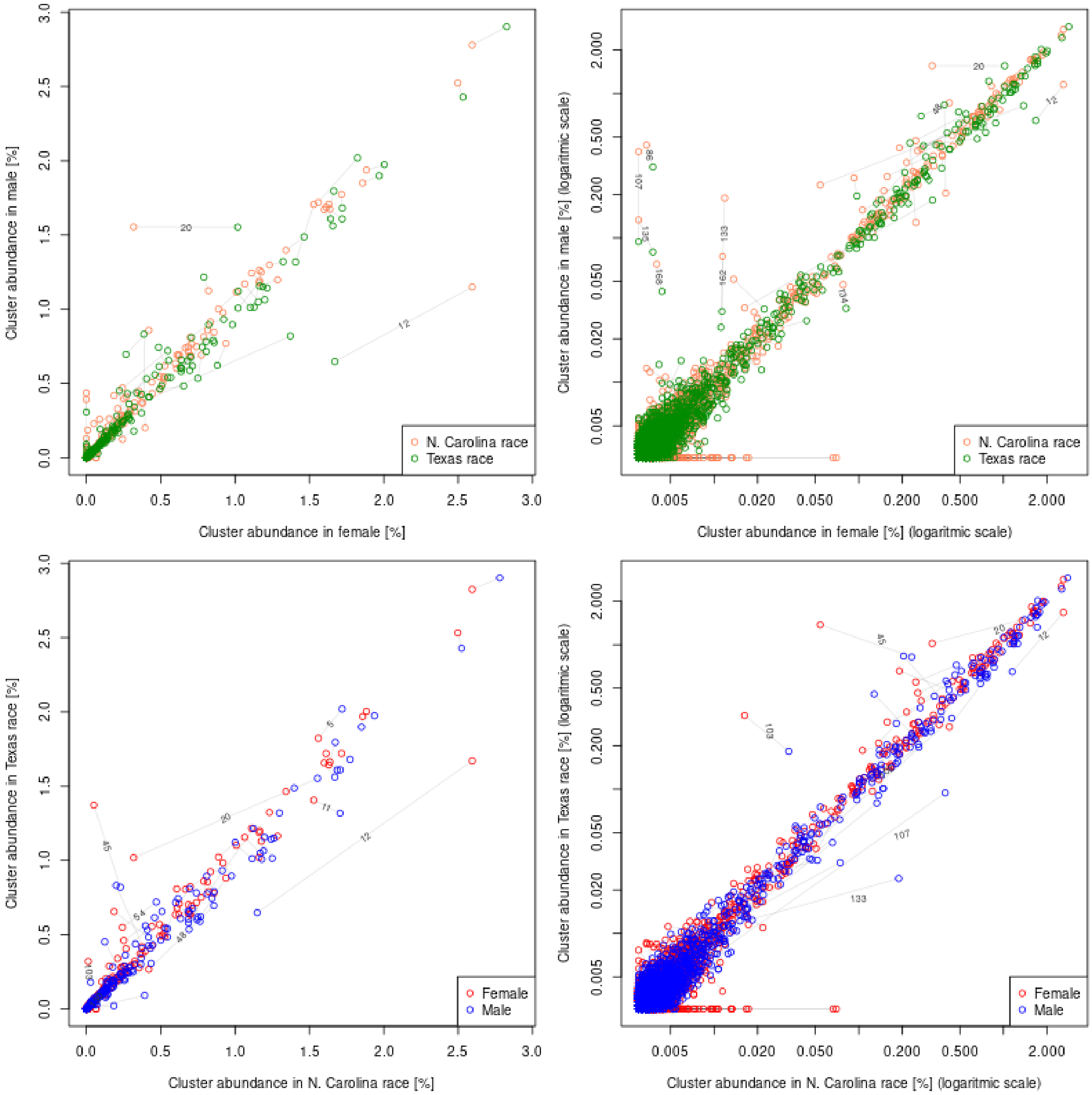

**Figure S3.** The localization of Cl12 (green), Cl134 5S (red-top) and Cl86 (red bottom) in XY and XYY males cytotypes. Metaphase chromosomes counterstained with DAPI (grey). Scale bar = 10 μm

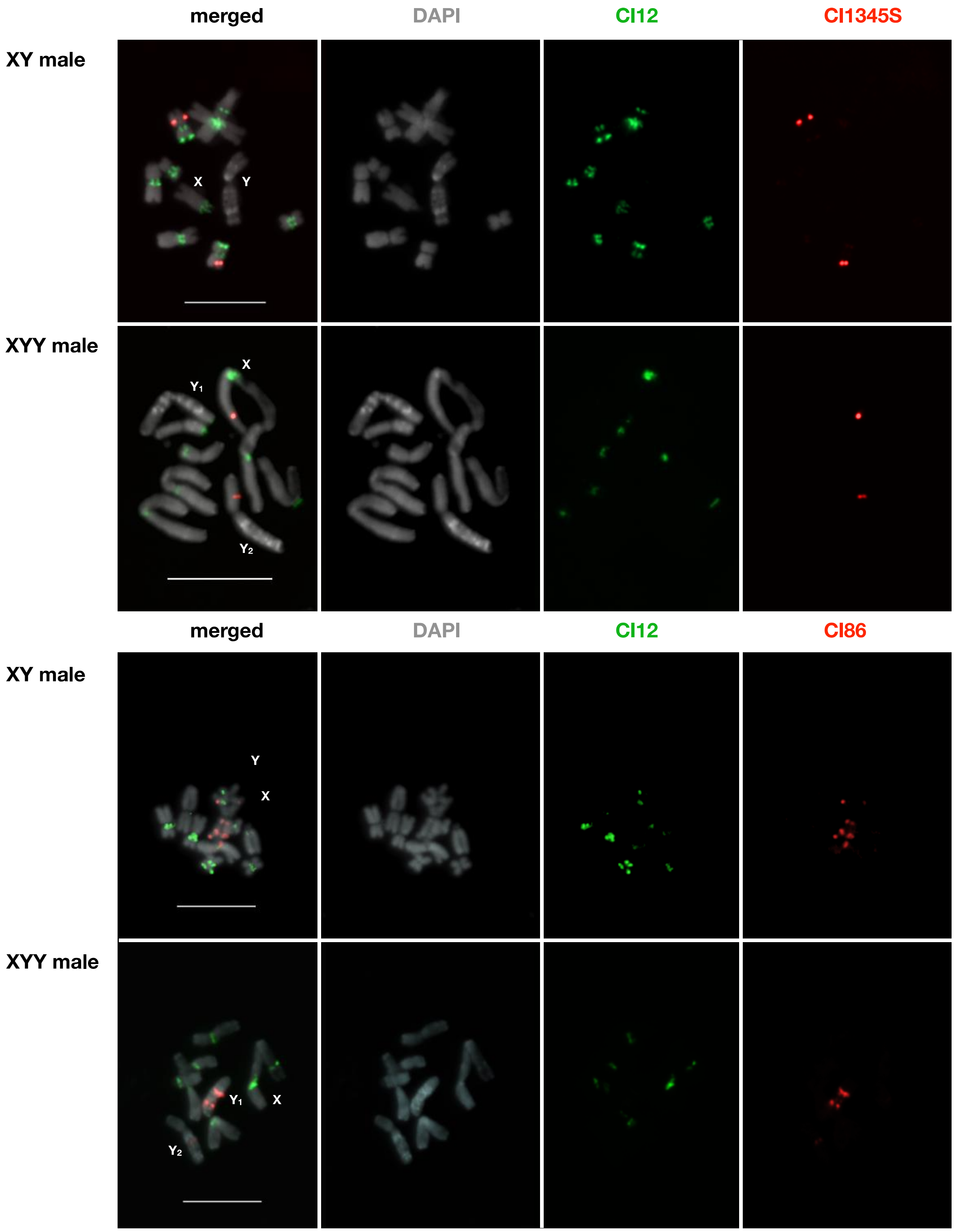

**Figure S4.** The localization of Cl12, Cl133 (red-top) and Cl135 (red bottom) in XY and XYY males cytotypes. Metaphase chromosomes counterstained with DAPI (grey). Scale bar = 10 μm

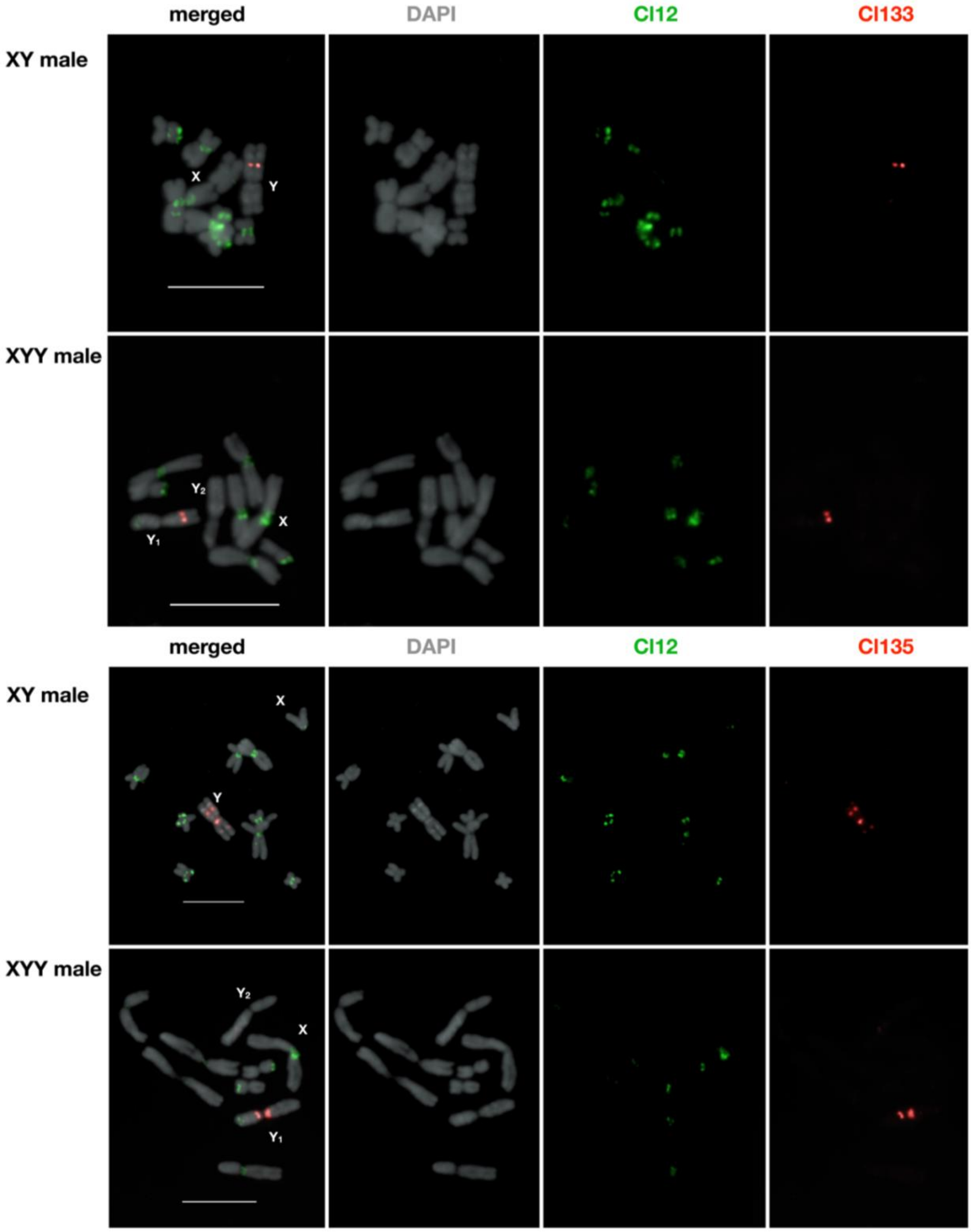

**Figure S5.** The localization of Cl12, Cl162 (red-top) and Cl168 (red bottom) in XY and XYY male cytotypes. Metaphase chromosomes counterstained with DAPI (grey). Scale bar = 10 μm

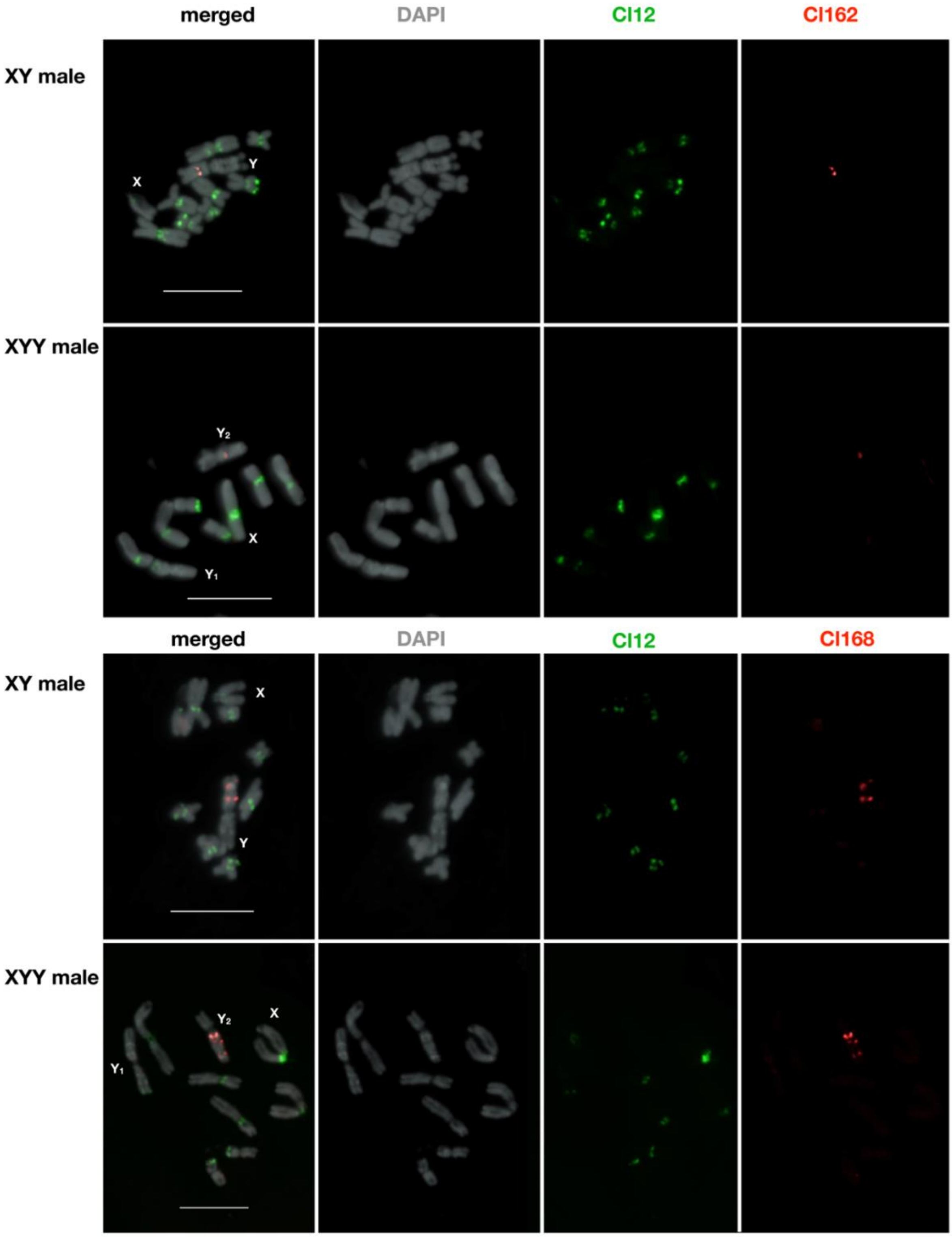

**Figure S6.** The scheme of satellite distribution in the XY and XYY male cytotypes of *R. hastatulus*. The satellite position along the chromosomal axis length is determined by ImageJ RGB plot diagram function, and its approximate order is shown above. Note different position of Cl12 between A4 in XY cytotype Texas and A4 in XYY cytotype North Carolina, confirming an autosomal inversion on this chromosomal pair.

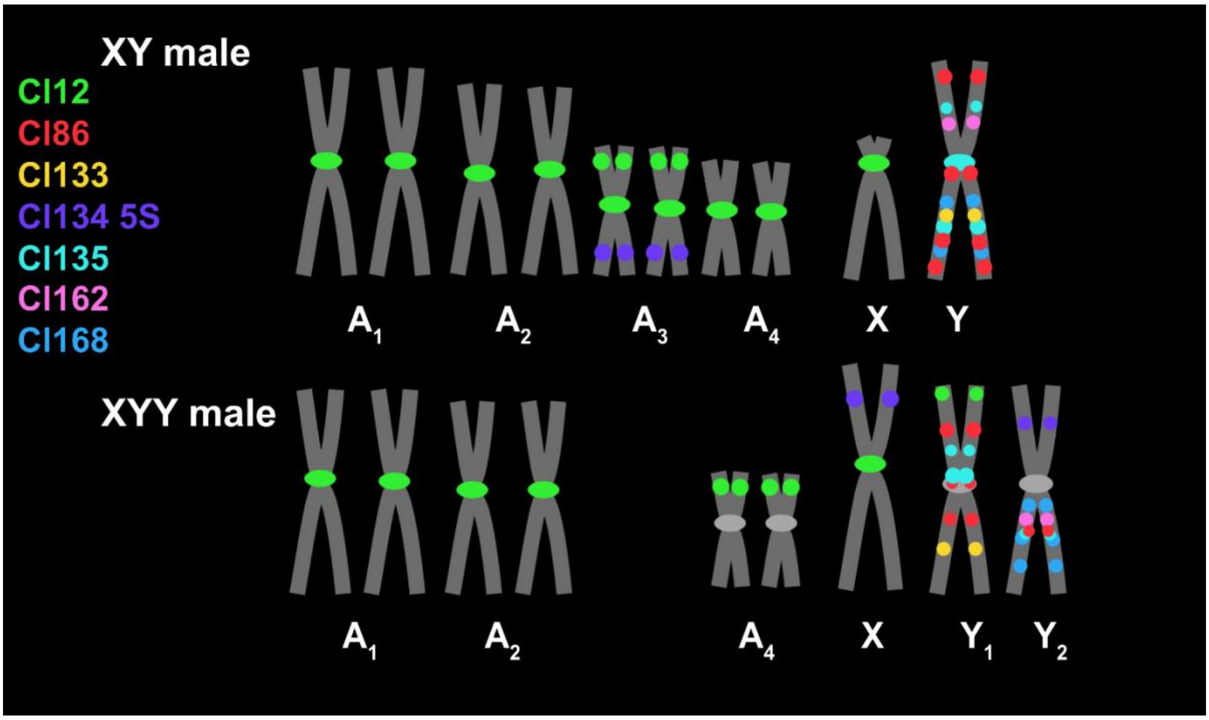

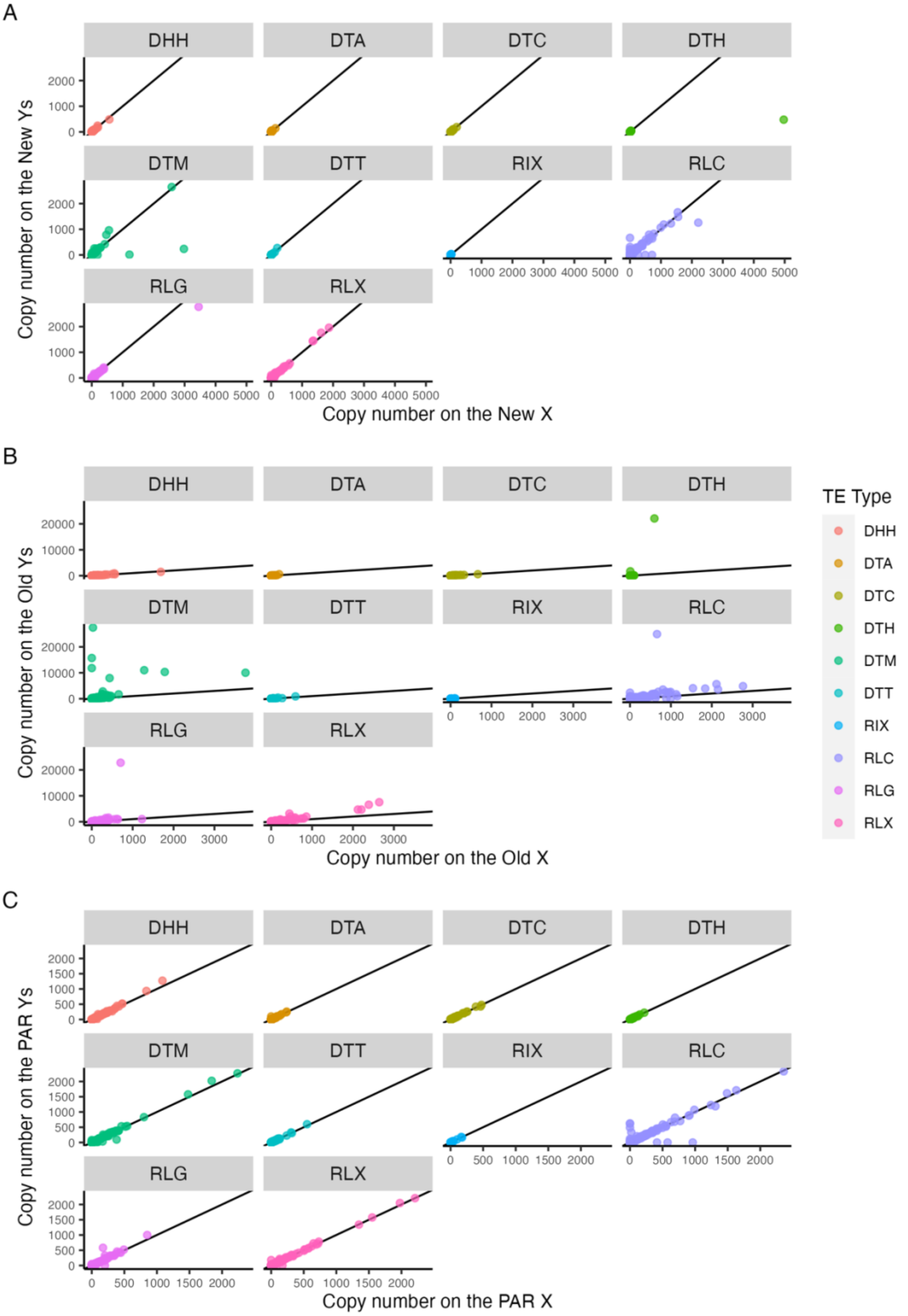


**Figure S7**. TE family copy number by TE Type. **A:** Comparison of TE copy number on the new sex-linked regions. **B:** Comparison of TE copy number on the old sex-linked regions. **C:** Comparison of TE copy number on the pseudo-autosomal regions (PARs). The three-letter transposon codes in the legend are from Wicker et al. 2007. DHH: helitron, DTA: hAT, DTC: CACTA, DTH: Harbinger, DTM: Mutator, DTT: Tc1-Mariner, RIX: unknown LINE (long interspersed nuclear element), RLC: Copia, RLG: Ty3, RLX: unknown LTR (long terminal repeat)

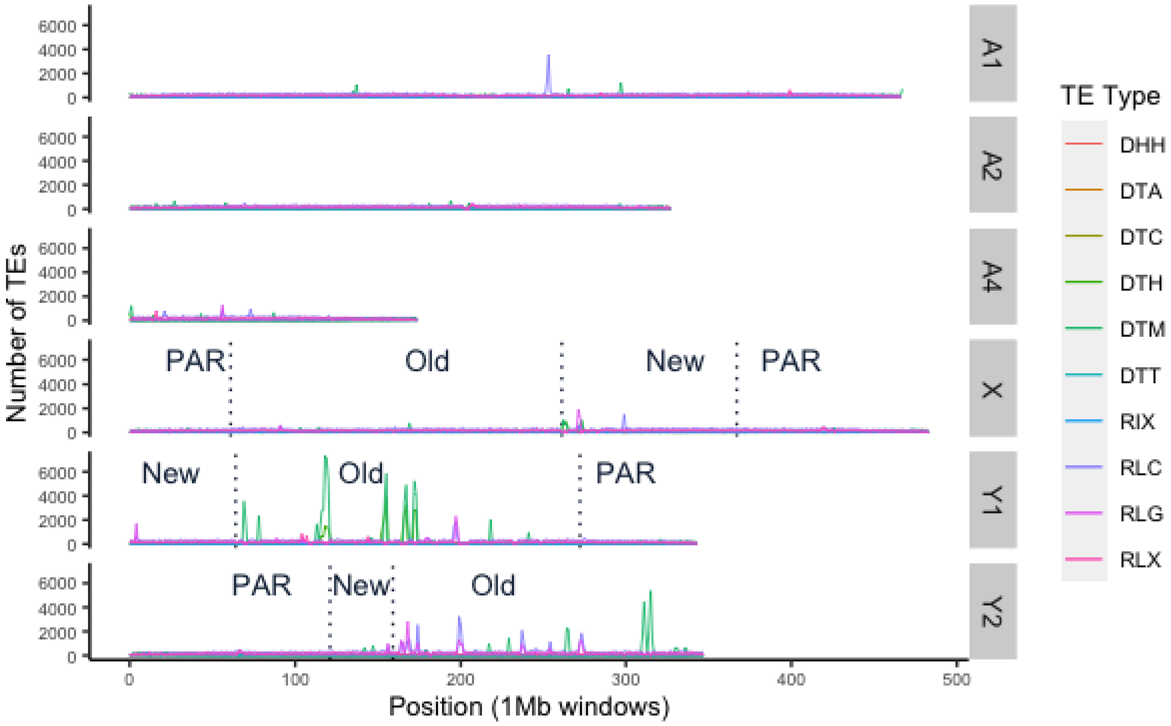


**Figure S8**. TE count landscape. Number of TEs per 1Mb window, coloured by TE type.
